# Temporal Chunking as a Mechanism for Unsupervised Learning of Task-Sets

**DOI:** 10.1101/713156

**Authors:** Flora Bouchacourt, Stefano Palminteri, Etienne Koechlin, Srdjan Ostojic

## Abstract

Depending on environmental demands, humans can learn and exploit multiple concurrent sets of stimulus-response associations. Mechanisms underlying the learning of such task-sets remain unknown. Here we investigate the hypothesis that task-set learning relies on unsupervised chunking of stimulus-response associations that occur in temporal proximity. We examine behavioral and neural data from a task-set learning experiment using a network model. We first show that task-set learning can be achieved provided the timescale of chunking is slower than the timescale of stimulus-response learning. Fitting the model to behavioral data confirmed this expectation and led to specific predictions linking chunking and task-set retrieval that were borne out by behavioral performance and reaction times. Comparing the model activity with BOLD signal allowed us to identify neural correlates of task-set retrieval in a functional network involving ventral and dorsal prefrontal cortex, with the dorsal system preferentially engaged when retrievals are used to improve performance.

## Introduction

Synaptic plasticity is believed to constitute the neurobiological basis of learning and memory. Changes of synaptic strength based on the activity of pre- and post-synaptic neurons were first postulated by Hebb [Hebb, 1949] and later confirmed in electrophysiological experiments [Frégnac et al., 1988; Levy and Steward, 1983; Lisman, 2003; Lynch et al., 1977; Malenka and Nicoll, 1999]. Such synaptic changes in turn modify the activity in the network as well as the response to incoming stimuli, and can implement stimulus-action learning [Bathellier et al., 2013; Fusi et al., 2007; Reynolds et al., 2001; Rioult-Pedotti et al., 2000; Rumpel et al., 2005; Schultz and Dickinson, 2000; Xiong et al., 2015]. Understanding how synaptic changes at individual synapses are related to learning in behaving animals remains however a daunting challenge, especially for complex cognitive tasks that go beyond simple stimulus-response associations.

Depending on environmental demands, humans engaged in a given task are capable of learning and exploiting multiple concurrent strategies. For instance, in the classical Stroop task [MacLeod, 1991; Stroop, 1935], an identical stimulus like a colored word leads to different responses depending on whether the current requirement is to read the word or identify its color. Human subjects are able to learn to flexibly switch between these two different stimulus-response associations, often called task-sets [Sakai, 2008]. Studies of task-sets learning predominantly rely on abstract models that describe behavioral learning without any physiological constraint [Botvinick et al., 2009; Collins and Koechlin, 2012; Collins and Frank, 2013; Daw et al., 2005, 2011; Dayan and Daw, 2008; Franklin and Frank, 2018; Russek et al., 2017]. While these models are able to capture computational aspects of behavior, and correlate them with physiological measurements [Daw et al., 2011; Donoso et al., 2014; Koechlin and Hyafil, 2007; Niv, 2009; Wilson et al., 2014], understanding the underlying biophysical mechanisms is an open issue.

One hypothesis [Rigotti et al., 2010b] states that learning of task-sets, and more generally rule-based behavior, relies on unsupervised learning of temporal contiguity between events. Events that occur repeatedly after each other are automatically associated as demonstrated in classical conditioning experiments [Hawkins et al., 1983; Kahana, 1996; Rescorla, 1988; Rescorla et al., 1972; Sakai and Miyashita, 1991; Yakovlev et al., 1998]. If one thinks of individual stimulus-response associations as abstracted events, temporally chunking two or more such events effectively corresponds to learning a simple task-set or association rule. Hebbian synaptic plasticity naturally leads to unsupervised learning of temporal contiguity between events [Blumenfeld et al., 2006; Fusi, 2002; Fusi et al., 2007; Griniasty et al., 1993; Li and DiCarlo, 2010; Ostojic and Fusi, 2013; Preminger et al., 2009; Soltani and Wang, 2006; Wallis et al., 1993], and therefore provides a natural biological mechanism for learning task-sets [Rigotti et al., 2010b].

Here we explore the hypothesis that temporal contiguity is implemented in a neural network through simple Hebbian learning. Recurrence enables the network to learn from its own activity and thus synaptic links are created between events separated in time. If the network is composed of hierarchical layers of cells selective to mixtures of stimuli and responses, task-sets of increasing complexity can be encoded. We test this hypothesis in the context of human decision-making. To this end, we consider a specific experimental task [Collins and Koechlin, 2012] where subjects have to learn multiple task-sets by merging associations between a unique set of stimuli and responses. We show that in order to encode these concurrent task-sets efficiently, plasticity between cells selective to a mixture of both a stimulus and a response has to be slower than plasticity between cells selective to either a stimulus or a response.

Fitting this model to behavior allows us to make specific predictions based on the hypothesis that task-set learning relies on temporal chunking of events. One prediction pertains to the case when a task-set is retrieved correctly, an another one to the case when this retrieval is maladaptive. We show that these predictions are borne-out by the behavioral data at the level of individual subjects. Moreover, we show that the time-series of the inference signal predicting task-set retrieval in the model correlates with BOLD signal recorded from fMRI [Donoso et al., 2014] in a functional network engaging medial and dorsal prefrontal cortex. Dorsomedial and dorsolateral prefrontal cortex are engaged specifically when the retrieval of a task-set is used for optimal performance. On the contrary, ventromedial prefrontal cortex is engaged negatively, irrespective of a possible task-set retrieval. This is equivalent to tracking the compatibility, when a reward is received, between the network layer encoding task-sets, and the network layer encoding one-to-one stimulus-response associations.

These results show that simple Hebbian mechanisms and temporal contiguity may parsimoniously explain the learning of complex, rule-based behavior.

## Results

### Behavioral evidence for task-set-driven behavior

To investigate the mechanisms for task-set learning, we examined a specific experiment performed by 22 human subjects (Experiment 1 [Collins and Koechlin, 2012], see Materials and Methods). In each trial, the subjects had to associate a visual stimulus with a motor response (Fig. 1a). The subjects needed to learn the correct associations based on a feedback signal, which was misleading in 10% of the trials (see Methods). The set of correct stimulus-response associations, which we will denote as *task-set* in the following, was fixed during a block of trials of random length (called an *episode*), and changed repeatedly to a non-overlapping set without explicit indication. As the feedback was not fully reliable, the subjects could not directly infer the task-set changes from the feedback on a single trial, but needed to integrate information.

**Figure 1:**
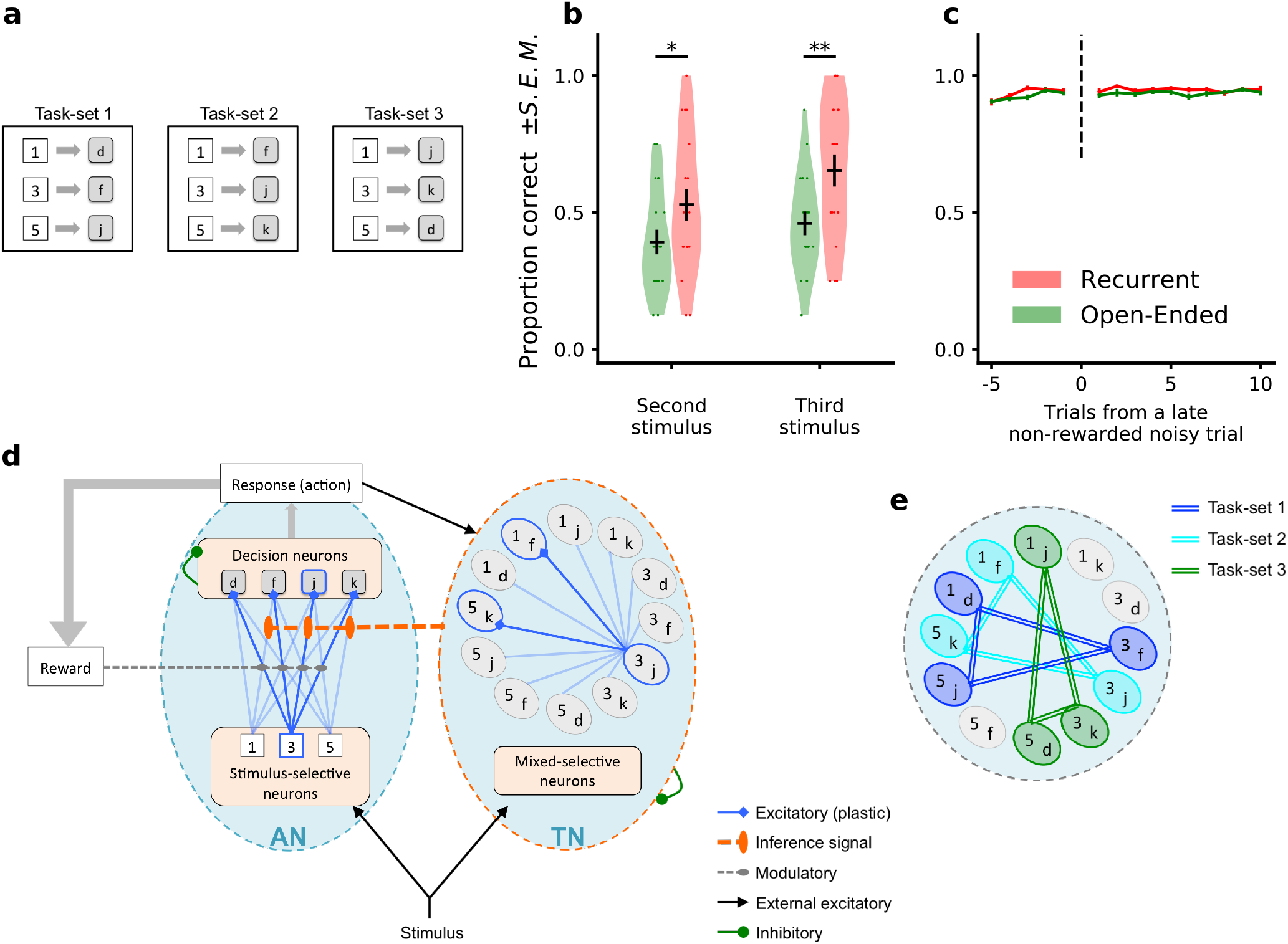
Task-set learning experiment and subject behavior. **a**, Schematic of the behavioral task. Subjects had to learn associations between visual stimuli (represented here as 1, 3, 5) and motor responses (represented here as *d, f, j, k*). The set of correct stimulus-response associations, denoted as *task-set*, was fixed during a block of trials of random length. The schematic shows the 3 task-sets used in the recurrent session. The task-sets are non-overlapping from one episode to another in both the recurrent and the open-ended session, meaning that an episode switch produces a change of correct responses for all stimuli. **b**, Proportion of correct responses to stimuli seen for the first time after the first correct response in an episode, during the last third of each experimental session. These newly seen stimuli are labeled *second* or *third* according to their order of appearance. Dots display the average for each subject. Violin plots display the shape of each distribution over subjects (Scott’s rule). The black lines outline the mean ± s.e.m. **c**, Performance preceding and following a trial with misleading feedback (non-rewarded correct response), at the end of episodes, averaged over all subjects (±s.e.m.). The subjects performance did not change after a misleading feedback if it occurs at the end of an episode, after being trained on the current task-set. **d**, Illustration of the network model. The associative network (AN) is composed of a set of stimulus-selective populations and a set of action-selective populations. The synaptic weights between the two sets of populations are modified through a reward-modulated, activity-dependent Hebbian plasticity rule. At each trial, an action is selected via a soft and noisy winner-take-all mechanism with respect to the current set of synaptic weights. The task-set network (TN) is composed of neural populations selective to conjunctions of one stimulus and one action. Its activity is driven by the associative network’s activity and by its own recurrent connectivity. The sequential activation of neural populations in the task-set network induces the potentiation of the synapses between them. An inference signal from the task-set network to the associative network biases future behavior according to the patterns of activation in the task-set network. **e**, Illustration of the perfect, fully chunked encoding in the task-set network of the 3 non-overlapping task-sets from the recurrent session.

The subjects’ behavior was compared between an *open-ended session*, in which the valid task-set was different in each episode, and a *recurrent session* in which only three task-sets appeared repeatedly. In the open-ended session, as each task-set was seen only once, a correct response to one stimulus bore only minimal information about the correct responses to the other stimuli (the responses to the three stimuli had to be different). In contrast, in the recurrent session, a correct response to a given stimulus fully predicted the correct responses to the other stimuli. Learning full task-sets rather than individual associations therefore allowed subjects to increase their performance.

Behavioral data indicated that beyond individual stimulus-response associations, subjects indeed learned task-sets in the recurrent session [Collins and Koechlin, 2012]. Additional evidence in that direction is displayed in Fig. 1b, where we show the proportion of correct responses to stimuli seen for the first time after the first correct response in an episode. This quantity was determined for the last third of the session, when the subjects had already experienced several times the three re-occurring task-sets within the recurrent session. If the subjects had perfectly learned the task-sets, they could directly infer the correct response to the newly seen stimulus from a single correct response to another stimulus. The data shows that indeed some subjects perfectly learned full task-sets, so that their performance is maximal after the first correct trial. The performance averaged over all subjects was significantly higher in the recurrent session compared to the open-ended session (T-test on related samples, second stimulus: 0.53 ± 0.05 vs 0.39 ± 0.04, t=2.1, p=0.049; third stimulus: 0.65 ± 0.05 vs 0.46 ± 0.03, t=3.1, p=0.0049), demonstrating that subjects exploited information from the correct response to a given stimulus to infer the correct response to other stimuli. An important variability was however observed among subjects, as most of them did not learn task-sets perfectly, and some not at all (a point we return to later).

An additional observation consistent with task-set learning was that subjects do not modify their behavior following a misleading noisy feedback occurring late in an episode (Fig. 1c, recurrent session: 0.94 ± 0.01 before, 0.94 ± 0.01 after, t=0.25, p=0.80; open-ended session 0.94±0.01 before, 0.93±0.01 after, t=0.68, p=0.50). An isolated misleading negative feedback after extensive learning in an episode should be ignored because inconsistent with the current task-set. A switch to another task-set or simply a change in a single stimulus-response association would be detrimental to performance. This negative feedback is indeed ignored by the subjects, indicating again they learn sets rather than individual stimulus-response associations.

### A network model for learning task-sets by chunking stimulus-response pairs

To examine the hypothesis that task-set-driven behavior emerges from unsupervised chunking of stimulus-response pairs, we studied a neurally-inspired network model (Fig. 1d), that built on previous modeling studies of a trace conditioning task in monkeys [Fusi et al., 2007; Rigotti et al., 2010b]. The model consisted of two subnetworks, which we refer to as the Associative Network (AN) and the Task-set Network (TN). The associative network is a simplified version of a neural decision-making network [Fusi et al., 2007; Wang, 2002; Wong and Wang, 2006]. It consists of a set of stimulus-selective populations and a set of action-selective populations. The stimulus-action associations are learned through reward-modulated plasticity on the synapses between the two sets of populations [Fusi et al., 2007].

The task-set network consists of neural populations that display mixed-selectivity to conjunctions of stimuli and actions [Rigotti et al., 2010b, 2013]. For instance, if the associative network generates the action *A***_2_** in response to the stimulus *S***_1_**, the corresponding population *S***_1_***A***_2_** is activated in the task-set network. Such mixed-selectivity response can be implemented through random projections from the associative network to the task-set network [Lindsay et al., 2017; Rigotti et al., 2010a], which for simplicity we don’t explicitly include in the model. Synapses between neural populations in the task-set network undergo temporal Hebbian learning, i.e. they are modified based on the successions of stimulus-response pairs produced by the associative network [Ostojic and Fusi, 2013; Rigotti et al., 2010b]. If two stimulus-response pairs are produced often after each other, the synapses between the corresponding mixed-selectivity populations in the task-set network are potentiated. When this potentiation exceeds a threshold set by recurrent inhibition, the two populations are chunked together and systematically coactivated when one of the two stimulus-response associations occurs. Thus by means of temporal chunking, this subnetwork implements a task-set as a merged pattern of co-activated populations. This coactivation is communicated to the associative network, where it biases the stimulus-action associations at the next trial towards those encoded by the active populations in the task-set network. This effective *inference signal* helps the associative network determine the correct response to a stimulus different from the one in the current trial, and therefore implements task-set retrieval in the network model. To keep the model easy to fit and analyze, this inference signal is implemented in a simplified manner, by directly modifying the synaptic weights in the associative network (see Discussion for more realistic physiological implementations). The strength of this modification is a parameter in the model that represents the strength of task-set retrieval (if it is zero, there is no retrieval). We will show that this specific parameter plays a key role in accounting for the variability in the subjects behavior.

### Task-set encoding in the network model enables task-set driven behavior

The task-set network is in principle able to chunk together stimulus-response pairs that occur often after each other. We first show how it enables task-set-driven behavior. Consider an idealized situation at the end of the recurrent session of the experiment where full chunking has taken place, and the pattern of connectivity in the task-set network directly represents the three task-sets (Fig. 1e). Due to the inference signal from the task-set network to the associative network, this pattern of connectivity will directly influence the responses to incoming stimuli.

The impact of this inference signal is the strongest at an episode change, when the correct set of stimulus-response associations suddenly shifts (Fig. 2a,e,i). The associative network always needs to discover the first correct association progressively by trial and error, by first depressing the set of stimulus-response synapses in the associative network corresponding to the previous task-set, and then progressively potentiating the synapses corresponding the new set of associations (Fig. 2a). In the absence of task-set inference, this learning process happens independently for each stimulus (Fig. 2b,c). In the presence of the idealized task-set network described above, once the first correct response is made, the task-set network produces the inference signal allowing the associative network to immediately recover the other two correct associations in the new episode (Fig. 2j,k). As a consequence, the overall performance is increased (Fig. 2e) due to a sudden increase in performance following the first correct response (Fig. 2f,g).

**Figure 2:**
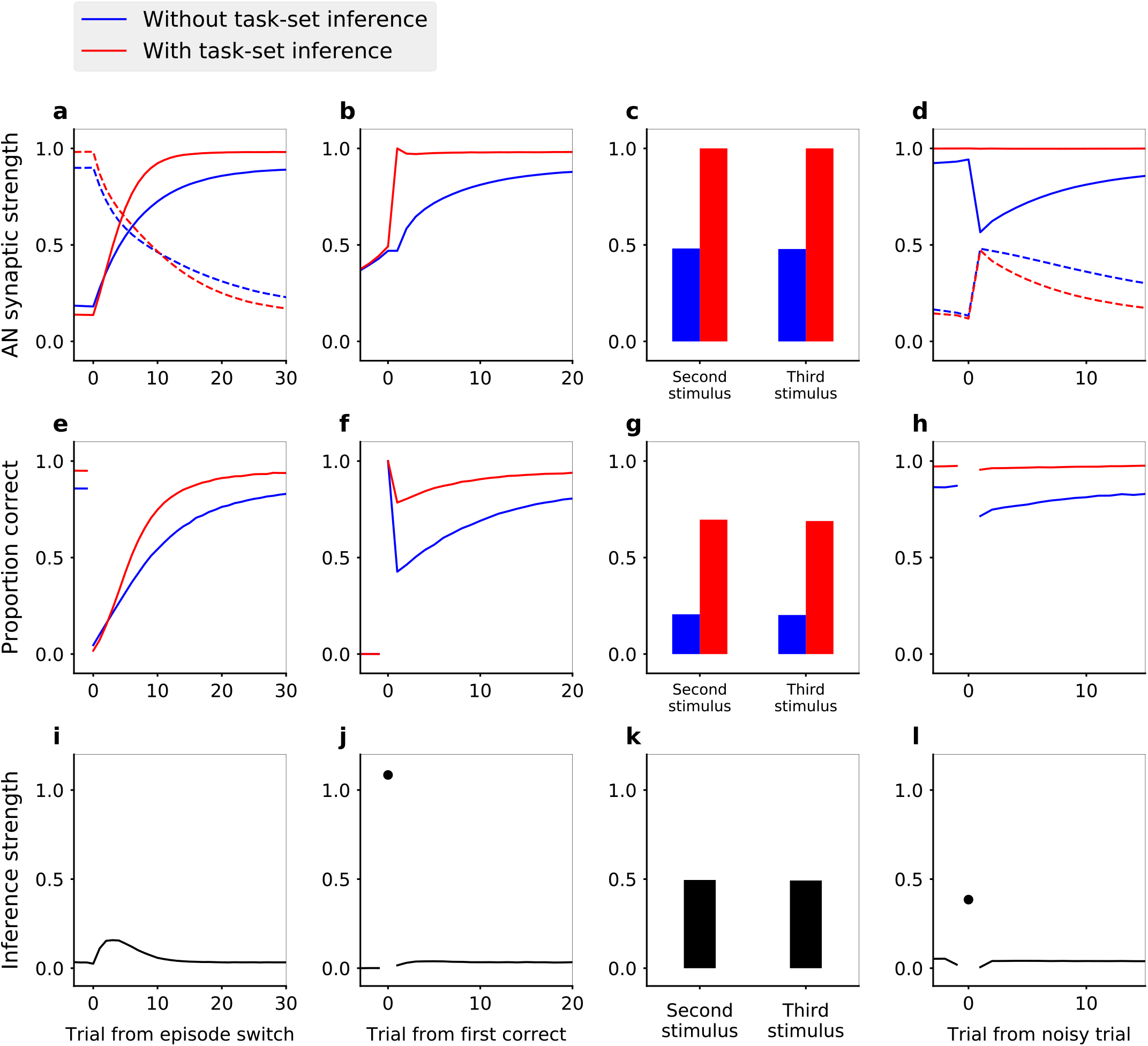
Task-set driven behavior in the network model with an idealized, perfect encoding of task-sets. The behavior of the model is compared in presence (red lines) and in absence (blue lines) of the inference signal from the task-set network, that allows task-set retrieval. **a,e,i**, Model dynamics following an episode switch (at trial zero, the correct task-set shifts without explicit indication). **a**, Strengths of synapses in the associative network between neural populations representing the new task-set (solid lines) and the previous task-set (dashed lines). **e**, Performance (proportion of correct responses) following the episode switch. **i**, Mean inference signal strength < *J*_*INC*_ · (1 − *J*^*AN*^) > from the task-set network to the associative network. **b,f,j**, Task-set retrieval: same quantities as in (a,e,i) for the stimulus seen for the first time after the first correct response in the new episode, aligned on the first correct response. **c,g,k**, Same quantities as in (a,e,i), for the two stimuli seen for the first time after the first correct response in an episode. As in Fig. 1b, These newly seen stimuli are labeled *second* or *third* relatively to their order of appearance. **d,h,l**, Effect of misleading feedback: same quantities as in (a,e,i,), aligned on a misleadingly non-rewarded correct trial at the end of episodes. Average of 5000 sessions of 25 episodes, with 10% of noisy trials. Network parameter values: *α* = 0.4, β = 7, *∊* = 0, *J*_*INC*_ = 1.

A second situation in which the task-set inference strongly manifests itself is the case of noisy, misleading feedback late in an episode. At that point, the associative network has fully learned the correct set of stimulus-response associations, and the performance has reached asymptotic levels. The network therefore produces mainly correct responses, but as on 10% of trials the feedback is misleading, it still occasionally receives negative reinforcement. In the absence of the task-set network, this negative feedback necessarily depresses the synapses that correspond to correct associations, leading to a decrease in performance on the following trials (Fig. 2d,h). In contrast, in the presence of the idealized task-set network, the inference signal that biases the behavior towards the correct task-set is present despite the occasional negative feedback, and therefore allows the network to ignore it (Fig. 2d,h,l). The encoding of task-sets in the task-set network pattern of connectivity therefore prevents the transient drop in performance, as seen in the experimental data (Fig. 1c).

### Speed-accuracy trade-off for learning task-sets in the network model

The idealized encoding described above requires that the task-set network effectively and autonomously learns the correct pattern of connections corresponding to the actual task-sets. We therefore next examined under which conditions synaptic plasticity in the task-set network leads to correct learning, i.e. correct temporal chunking.

Fig. 3a,c,e,g shows a simulation for a parameter set for which learning of task-sets proceeds successfully. At the beginning of the session, all populations within the task-set network are independent as all synaptic weights are below threshold for chunking. As the associative network starts producing correct responses by trial and error, the weights in the task-set network that connect correct stimulus-response pairs get progressively potentiated. While a fraction of them crosses threshold and leads to chunking during the first episode (and therefore starts producing the task-set inference signal), the majority does not, reflecting the fact that the first task-set is not fully learned at the end of the first episode. The potentiation in the task-set network continues over several episodes, and the weights in the task-set network that correspond to co-occurring stimulus-response pairs eventually saturate to an equilibrium value. This equilibrium value is an increasing function of the probability that two stimulus-response pairs follow each other, and of the potentiation rate in the task-set network (see Methods). The equilibrium synaptic weights in the task-set network therefore directly reflect the temporal contiguity between stimulus-response pairs [Ostojic and Fusi, 2013] and thus encode the task-sets. If the equilibrium value is larger than the inhibition threshold in the task-set network, this encoding will lead to the chunking of the activity of different populations and generate the inference signal from the task-set network to the associative network. The inference sets in progressively as the synaptic plasticity in the task-set network advances, and increasingly biases the behavioral output produced by the associative network.

**Figure 3:**
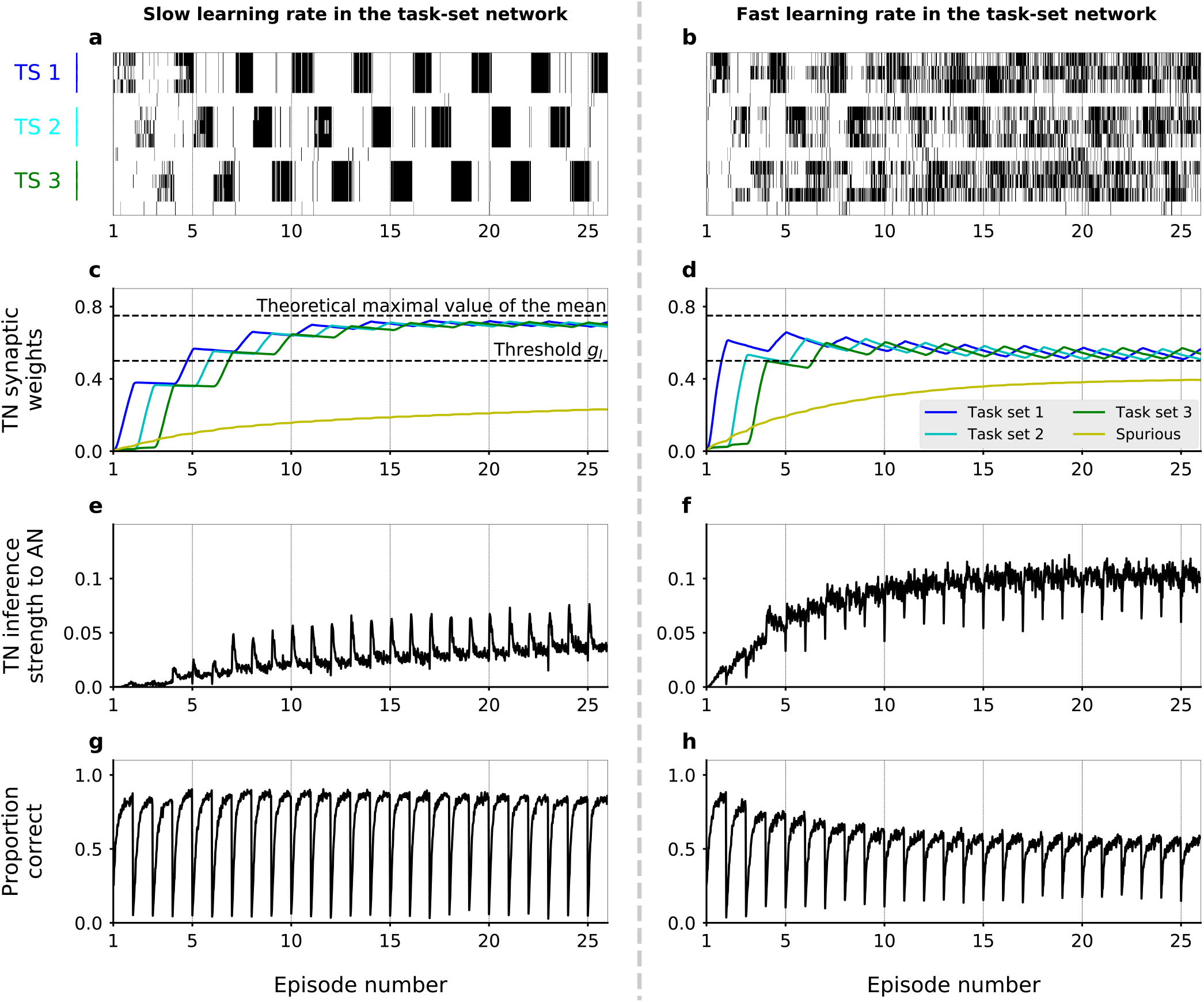
Dynamics of task-set learning. Left column: slow learning rate in the task-set network (TN) (*Q_P_* = 0.17); right column: fast learning rate in the task-set network (*Q_P_* = 0.4). **a,b**, Activation of neural populations in the task-set network as a function of time during one session. In (a), learning dynamics proceed correctly and lead to the chunking of populations that correspond to the same taskset. As a result, the activation of one stimulus-response association causes the co-activation of the other two in the same task-set. In contrast, in (b) learning does not proceed correctly and chunking does not take place. **c,d**, Average values of task-set network synaptic strengths between neural populations corresponding to each of the three correct task-sets, as well as “spurious” synaptic strengths between neural populations from different task-sets or that do not correspond to any task-set at all. **e,f**, Average value of the inference signal from the task-set network to the associative network connectivity. **g,h**, Performance of the network. Task-sets presentation is periodic for illustration purposes. (a,b) corresponds to 1 run of the recurrent session. (c,d,e,f,g,h) corresponds to the average over 500 runs of the recurrent session. The values of parameters other than *Q_p_* were *α* = 0.4, *β* = 7, *∊* = 0, and *J*_*INC*_ = 0.7.

Learning in the task-set network is however strongly susceptible to noise, and need not necessarily converge to the correct representation of task-sets. One important source of noise is the exploratory period following an episode switch, during which the associative network produces a large number of incorrect responses while searching for the correct one. If the potentiation rate in the task-set network is too high, the synaptic weights in the task-set network may track too fast the fluctuating and incorrect stimulus-response associations produced by the associative network, and quickly chunk together pairs of events that do not correspond to a correct task-set (Fig. 3b). Once these events are chunked together, the task-set network sends an incorrect inference signal to the associative network, and generates further incorrect associations (Fig. 3f,h). As the network learns in an unsupervised fashion from its own activity, this in turn leads to more incorrect associations in the task-set network. In such a situation, the presence of the task-set network is at best useless and at worse detrimental to the performance of the network as a whole.

To determine under which conditions the plasticity in the task-set network leads to the correct learning of task-sets, we systematically varied the associative and task-set networks learning rates and compared the performance in the models with and without the inference signal from the task-set network. Our results show that the presence of task-set inference improves the network performance when the task-set network learning rate is slower than the associative network learning rate (Fig. 4a). As illustrated in Fig. 3b,f,h, when learning in the task-set network is too fast, the network tracks noisy associations produced by the associative network, because of noise in the experimental feedback or because of errors made at the transition between episodes. In contrast, slow learning allow the task-set network to integrate information over a longer timescale. While in principle it would be advantageous to learn the task-set structure as quickly as possible, the requirement to average-out fluctuations due to erroneous feedback sets an upper-bound on the learning rate in the task-set network. This is an instance of the classical speed-accuracy trade-off.

**Figure 4:**
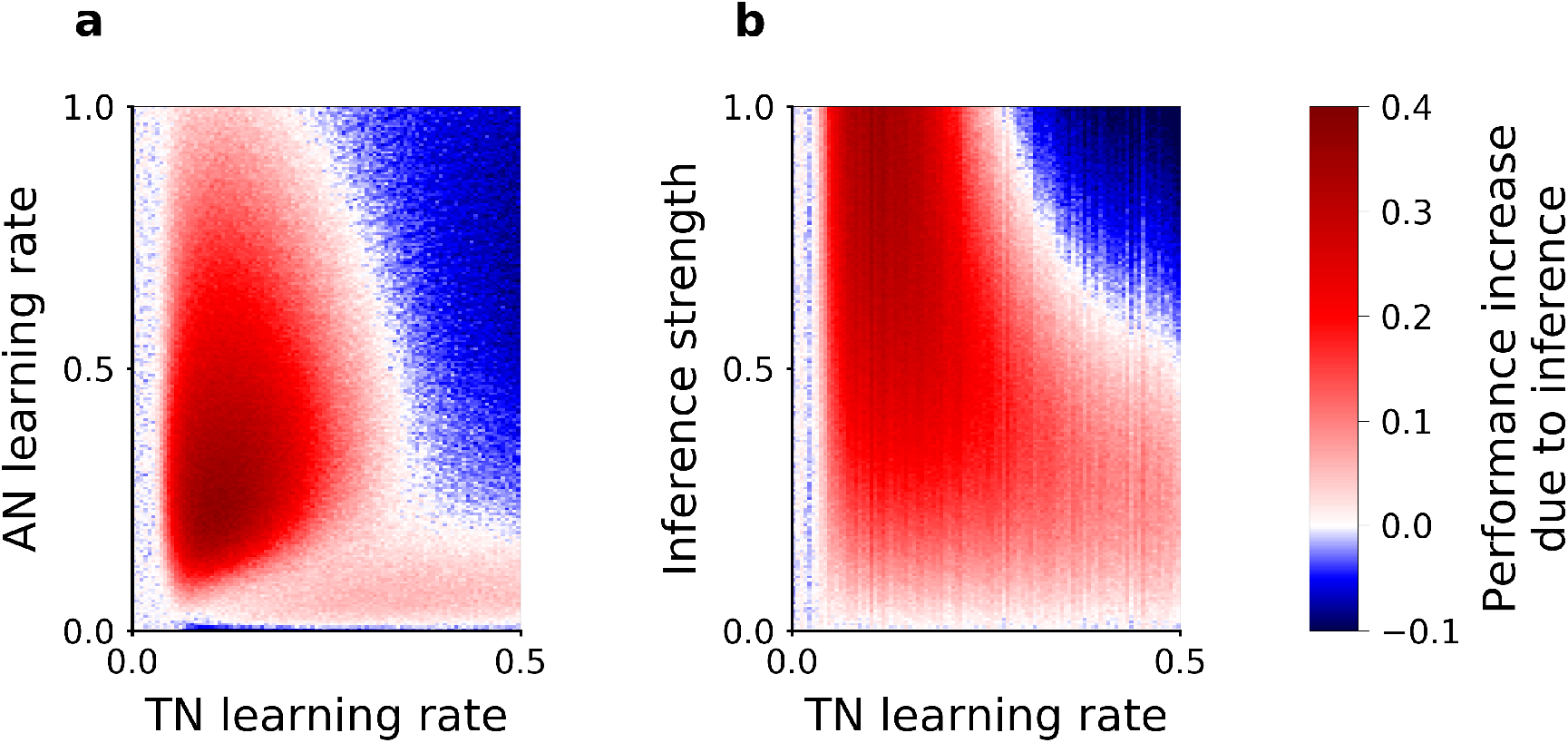
Slow versus fast learning: conditions for correct encoding of task-sets in the network model. **a**, Difference in the performance of the network model with or without task-set inference, plotted as function of the associative network learning rate *α* and the task-set network learning rate *Q****_P_***, (with *β* = 7 and inference strength *J*_*INC*_ = 0.7). **b**, Same difference in performance but plotted as function of the inference strength *J*_*INC*_ and the task-set network learning rate *Q****_P_***, (with *β* = 7 and associative network learning rate *α* = 0.4). We computed the performance averaged over the 5 first correct responses for a stimulus, in the last third of the session, on an average of 200 runs of the recurrent session and with 10% noisy trials.

The correct learning of task-sets also depends on the strength of the inference signal. While strong inference leads to strong task-set retrieval and potentially large performance improvement, it also makes the network more sensitive to incorrect chunking in the task-set network. Our simulation show that larger inference strengths need to be compensated for by lower learning rates in the task-set network to produce an improvement in the performance (Fig. 4b). This is another manifestation of the speed-accuracy trade-off.

### Fitting the model to behavioral data

Having described the dynamics in the model, we next proceeded to fit the model parameters to the subjects’ behavioral data. The full network model contains only five free parameters, which we determined independently for each subject by maximizing the likelihood of producing the same sequence of responses. To determine the importance of task-set retrieval in the model, we compared the fit obtained from our two nested models: the full model (associative network connected to the task-set network, 5 parameters), as well as the associative network model alone, without the inference signal that allows task-set recovery (3 free parameters). The full model with task-set inference provides a significantly better fit to the behavioral data in the recurrent session than does the model without task-set inference (Fig. 5a, Bayesian Information Criterion (BIC), T-test on related samples *t* = 14, *p* = 4.3 *·* 10^−12^). In the open-ended session, in which a given task-set never reoccurs between episodes, the two models provide instead indistinguishable fits. On average over subjects, the learning rate in the associative network (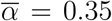, *σ* = 0.0073) is twice the learning rate in the task-set network (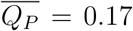, *σ* = 0.0070), which is consistent with our initial prediction that the learning rate in the task-set network needs to be slower than in the associative network.

**Figure 5:**
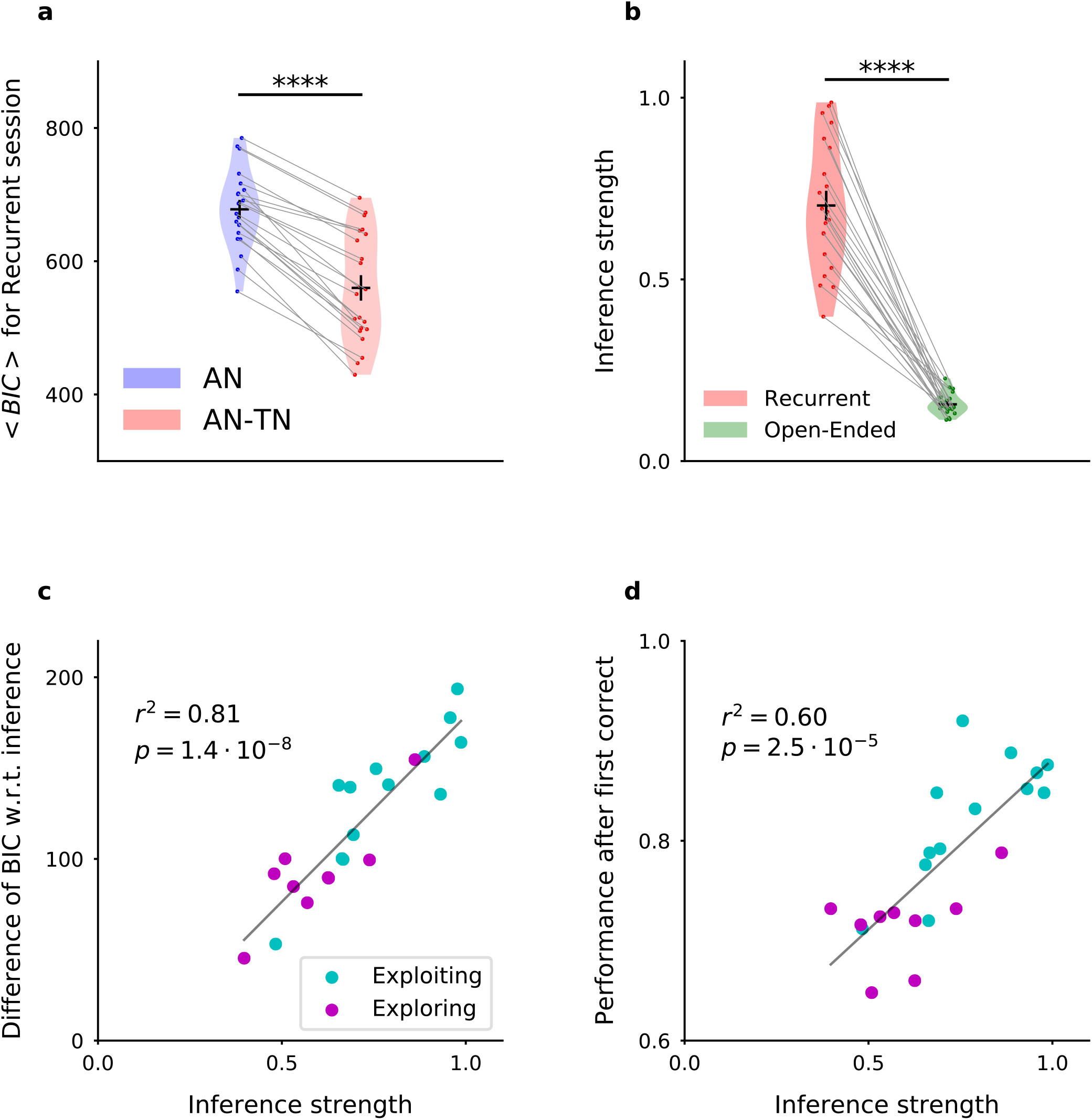
Fitting the model to experimental data: the model with inference (AN-TN) captures the statistical structure of the data, and accounts for the variability between subjects. **a**, Model comparison for the recurrent session. Bayesian Information Criterion, computed as in [Bishop, 2007], for the models with and without task-set inference. The model provides a significantly better fit with inference than without. **b**, Estimate of the inference strength *J*_*INC*_ from the task-set network to the associative network connectivity in the model with task-set inference, for both sessions. **c**, Subject by subject difference between BIC values obtained for models with and without task-set inference, as a function of the inference strength parameter. Subjects are classified as “exploiting” or “exploring” from a post-test debriefing. The grey line displays a least-squares regression. **d**, Subject by subject performance following the first correct trial in an episode, as a function of the inference strength parameter. The performance was computed by considering the 10 trials following the first correct trial of each episode. The grey line displays a least-squares regression.

As mentioned earlier, an important behavioral variability was present among subjects. This variability was particularly apparent in the performance following an episode switch, where some subjects increased their performance much faster than others in the recurrent session, compared to the open-ended session (Fig. 1b). Inspecting the parameter values obtained for different subjects revealed that the most variable model parameter between the recurrent and the open-ended sessions was the strength of the inference signal for task-set retrieval in the model (Fig. 5b, T-test on related samples *t* = 14.8, *p* = 1.5 · 10^−12^). The value of this parameter appeared to directly account for the inter-subject variability, as it correlated with the difference between BIC values obtained for models with and without task-set inference (linear regression *r*^2^ = 0.81, *p* = 1.4 *·* 10^−8^, Fig. 5c) as well as with the subjects’ performance following the first correct trial in an episode (linear regression *r*^2^ = 0.60, *p* = 2.5 *·* 10^−5^, Fig. 5d). These findings further suggest the variability in that parameter is directly linked with the subject’s ability to recover task-sets. This was confirmed by examining the results of a behaviorally-independent post-test debriefing used in the original study to classify subjects as either exploiting task-set structure (“exploiting” subjects) or not (“exploring” subjects). Exploiting subjects systematically corresponded to higher performance on trials following a correct response, and higher values of the inference parameter in the model (Fig. 5c,d).

### Testing model predictions for task-set retrieval

We next examined a specific subset of experimental trials where task-set retrieval is expected to take place. In the model, how quickly two stimulus-response pairs are chunked together depends on how often they co-occur, as well as on the value of the learning rate in the task-set network. Once two pairs are chunked together, the correct response to the stimulus corresponding to one of the pairs leads to the retrieval of the task-set, and biases the response to the stimulus from the second pair. When the pairs are not chunked together, the responses to the two stimuli are instead independent. The basic prediction is therefore that the responses to the stimulus from the second pair should differ between trials when chunking has or has not taken place, depending on the learning progress.

We first tested this prediction in a situation where chunking should lead to the retrieval of the correct task-set. We focused on one trial in each episode, the trial that followed the first correct response (Fig. 6a). Running our model on the full sequence of preceding experimental events (on a subject-by-subject basis, using parameters fitted to each subject and actual sequences of stimuli and responses) produced a prediction for whether chunking had occurred for this trial (*chunked* or *independent*, Fig. 6a, orange and grey respectively). The model with inference predicted that the proportion of correct responses on chunked trials should be higher than on independent trials due to the inference signal implementing task-set retrieval. In the model without inference where the associative network is independent of the task-set network, the performance on the two types of trials is instead indistinguishable. Comparing the proportion of correct responses on experimental trials classified in the two categories showed a significant increase for chunked trials compared to independent trials (Fig. 6b: (i) model without inference *t* = 0.64, *p* = 0.53, (ii) model with inference *t* = 6.9, *p* = 1.4 *·* 10^−7^, (iii) subjects *t* = 2.8, *p* = 8.8 *·* 10^−3^, (iv) chunked trials, model without inference versus model with inference *t* = 11, *p* = 6.3*·*10^−10^, (v) chunked trials, model without inference versus subjects *t* = 5.9, *p* = 2.3 *·* 10^−5^), so that the model prediction with inference was directly borne out by experimental data. The task-set retrieval predicted by the model therefore led to a clear increase of subjects’ performance. Moreover, reaction times on chunked trials were significantly lower than on independent trials, showing that the inference helped subjects to be faster at responding (Fig. 6c, *t* = 8.7, *p* = 1.7 *·* 10^−9^). This provides a supplementary validation, as the model was not fitted on reaction times. Note that a potential confound could be induced if the chunked trials appeared on average later in an episode than independent trials. A direct comparison however showed that the distributions of chunked and independent trials were indistinguishable (Fig. S3a, *ks* = 0.085, *p* = 0.62).

**Figure 6:**
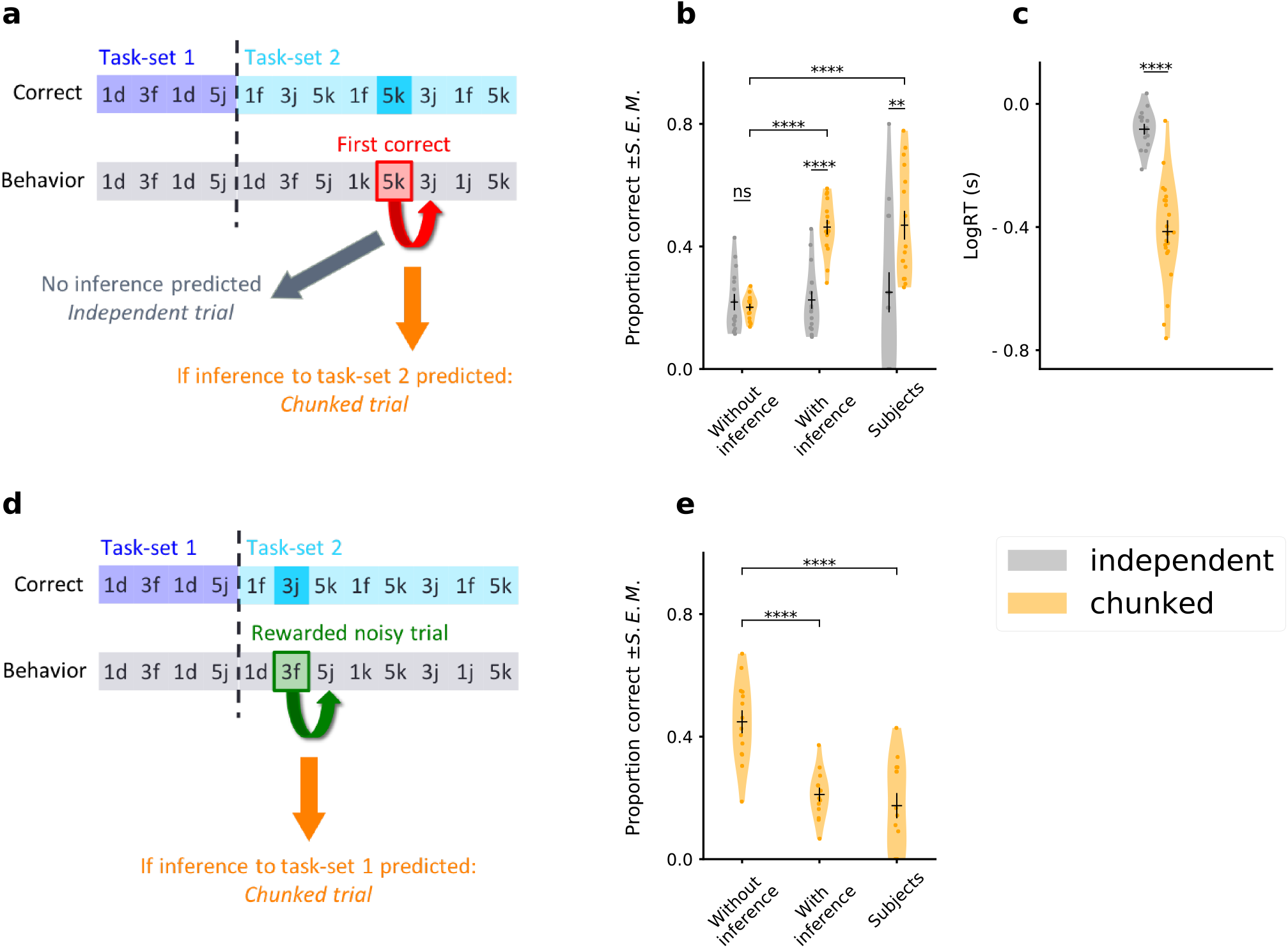
Testing the predictions of the temporal chunking mechanism on specific trials. **a**, Schematic of the prediction for correct task-set retrieval. For each episode, and subject by subject, we compute the probability of making a correct choice after the first correct trial following an episode switch. Trials are classified from a model-based criterium as “chunked” or “independent”, respectively depending on the presence or absence of an inference from the task-set network to the associative network. **b**, Because of task-set inference, the model predicts a significant increase of performance on chunked trials compared to independent trials. This is not predicted by the associative network alone (“Model without inference”). Subject’s performance on these trials matches the model with inference. The error bars are larger for the *independent* trials because this category contains half the number of data, as shown in figure S3. **c**, Log of subjects’ reaction times in seconds, for trials classified as chunked or independent. **d**, Schematic of the prediction for task-set retrieval following misleading rewarded trials. After each episode switch, the subject is making incorrect choices. On 10% of these trials the feedback is misleadingly rewarded (e.g. 3*f*, which corresponds to a correct association for the previous task-set, but not for the current task-set). Because of the inference from the task-set network, the previous task-set can be incorrectly inferred by the model from this positive feedback: it’s a “chunked” trial. **e**, Probability of making a correct association after a misleadingly rewarded noisy trial classified as a chunked trial by the model. Because of the inference from the task-set network, the model predicts an incorrect association at the next trial, producing a decrease in performance. This decrease is not predicted by the associative network alone. Subject’s performance on these trials matches the model with inference. Violin plots display the shape of each distribution (Scott’s rule). Dots display the average for each subject. The black lines outline the mean ± s.e.m.

We next tested the predictions of the model on trials where chunking leads to the retrieval of an incorrect task-set. Such a situation happens because of the presence of 10 % of trials with misleading feedback, which may indicate to the subject that the produced response was correct although it was not. Our model predicted that in this case incorrect task-set recovery leads to a decrease of the performance on the next trial. To test this prediction, we first detected the misleading trials, and then used the model to classify each of them as either chunked or independent (Fig. 6d). Comparing in the experimental data the responses on chunked trials with the performance of the model without task-set inference showed that indeed the subjects’ performance was significantly reduced when the model predicted an incorrect task-set retrieval (Fig. 6e: (i) chunked trials, model without inference versus model with inference *t* = 5.8, *p* = 1.0 *·* 10^−5^, (ii) chunked trials, model without inference versus subjects *t* = 5.2, *p* = 2.2 *·* 10^−5^).

The two behaviors described above (retrieval of a correct task-set after the first correct response, and retrieval of an incorrect task-set after a misleading feedback) cannot be predicted by the model without inference: we thus assessed the generative performance of our chunking mechanism and *falsified* the model without inference [Palminteri et al., 2017].

### Task-set inference related activity in ventromedial, dorsomedial, and dorsolateral prefrontal nodes

The neural populations in the task-set network represent neurons selective to conjunctions of stimuli and responses. Temporal synaptic chunking between these neural populations over the course of learning created task-sets in the model, and predicted precisely timed retrievals, correct or maladaptive, and borne out by individual subjects’ behavior. We aimed at understanding where this signal was implemented in the brain, using blood-oxygen-level-dependent signal recorded from functional magnetic resonance imaging (40 subjects, Experiment 2 [Donoso et al., 2014], see Fig. S2 for model fit on this dataset).

We first investigated neural correlates of the trial-by-trial synaptic strength of the chosen association in the associative network, *W*_*chosen*_, at the onset decision (controlling for trial difficulty, through time-series of reaction times, see Methods). As shown in Fig. S5 (top), we found positive linear effects in striatum and ventromedial prefrontal cortex, and negative linear effects in dorsal anterior cingulate cortex, anterior supplementary motor area and lateral prefrontal cortex, in both experimental sessions. These results confirm previous findings in the field of value-based decision making [Alexander and Brown, 2011; Chib et al., 2009; Daw et al., 2006; Donoso et al., 2014; Lebreton et al., 2009; Neubert et al., 2015; Palminteri et al., 2015; Tanaka et al., 2004].

We then focused on the correlations between BOLD signal and the trial-by-trial inference signal strength which we found to distinguish the two experimental sessions, computationally and behaviorally (at the onset feedback, trial difficulty and prediction error were controlled with *W*_*chosen*_ and the time series of positive rewards, see Methods). In the recurrent session (Fig. S5, bottom), BOLD activity correlated positively with the inference signal strength in dorsolateral prefrontal cortex, dorsal anterior cingulate cortex and anterior supplementary motor area; and negatively in ventromedial prefrontal cortex. In contrast, in the open-ended session, we found no significant positive or negative effect in frontal lobes corresponding to this parametric modulator. These results showed important differences between the two sessions, and suggested that a functional network engaging medial and dorsolateral prefrontal cortex is activated when the task-sets had to be learned and used (recurrent session versus open-ended session).

Hence we defined a functional network from significant BOLD activations in both sessions for the task-set network inference signal at the onset feedback. To do so, we performed a between-subjects ANOVA (contrasts REC+OE and -REC-OE, see Table 7 and Methods), which showed dorsomedial and dorsolateral prefrontal cortex correlated positively with the value of the inference signal. Ventromedial prefrontal cortex correlated negatively with the value of the inference signal, i.e. positively with the compatibility between encoding in the two subnetworks (see Methods) when a reward is received.

This functional network was used for a ROI analysis on the trial-by-trial inference signal to test the hypothesis of a specific effect of task-set retrieval in the recurrent session (ventromedial, dorsomedial and dorsolateral prefrontal cortex respectively left, middle, and right columns of Fig. 8a). These ROI were thus chosen from the analysis of a different contrast (REC+OE, see Methods) that did not promote differences (REC versus OE). First, we tested these ROIs at the onset feedback for *W*_*chosen*_ (top row) and for receiving positive reward (thus on “prediction” error, middle row): both were significantly encoded in the three ROIs, but no difference between experimental sessions was observed. The task-set network inference signal was also significantly encoded in the three ROIs (bottom row). However, the correlation with this signal in dorsomedial and dorsolateral prefrontal cortex was significantly stronger in the recurrent session, compared to the open-session (dmPFC: *t* = 3.0, *p* = 3.2 *·* 10^−3^; and dlPFC: *t* = 4.6, *p* = 1.6 *·* 10^−5^), i.e. when this signal was needed to improve performance in the task. Activations in ventromedial prefrontal cortex did not discriminate significantly between the two sessions (*t* = 0.16, *p* = 0.87). This analysis showed a difference of neural activity in the dorsal system corresponding to the necessity of learning and using (recurrent session) or not (open-ended session) the model of the task. We controlled our neural findings by performing analyses with ROI selected from several meta-analyses ([Glasser et al., 2016; Lancaster, 1997; Shirer et al., 2012; Yarkoni et al., 2011], see Methods). As shown in Fig. S6, dorsolateral and dorsomedial prefrontal cortex were specifically recruited in the recurrent session and implicated in the computation of the inference signal for task-set retrieval.

**Figure 7:**
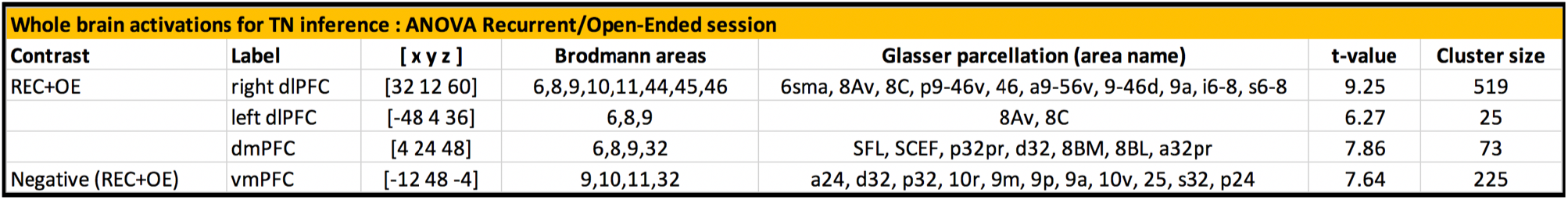
One-way ANOVA defining the functional network engaged in the TN inference signal computation. The functional network is defined from activations from the parametric modulator corresponding to the TN inference signal, in both sessions (contrasts REC+OE and -REC-OE, FWE *p* = 0.05). dlPFC: dorsolateral prefrontal cortex; dmPFC: dorsomedial prefrontal cortex; vmPFC: ventromedial prefrontal cortex; [x y z] are MNI coordinates; REC: Recurrent session; OE: Open-Ended session.

**Figure 8:**
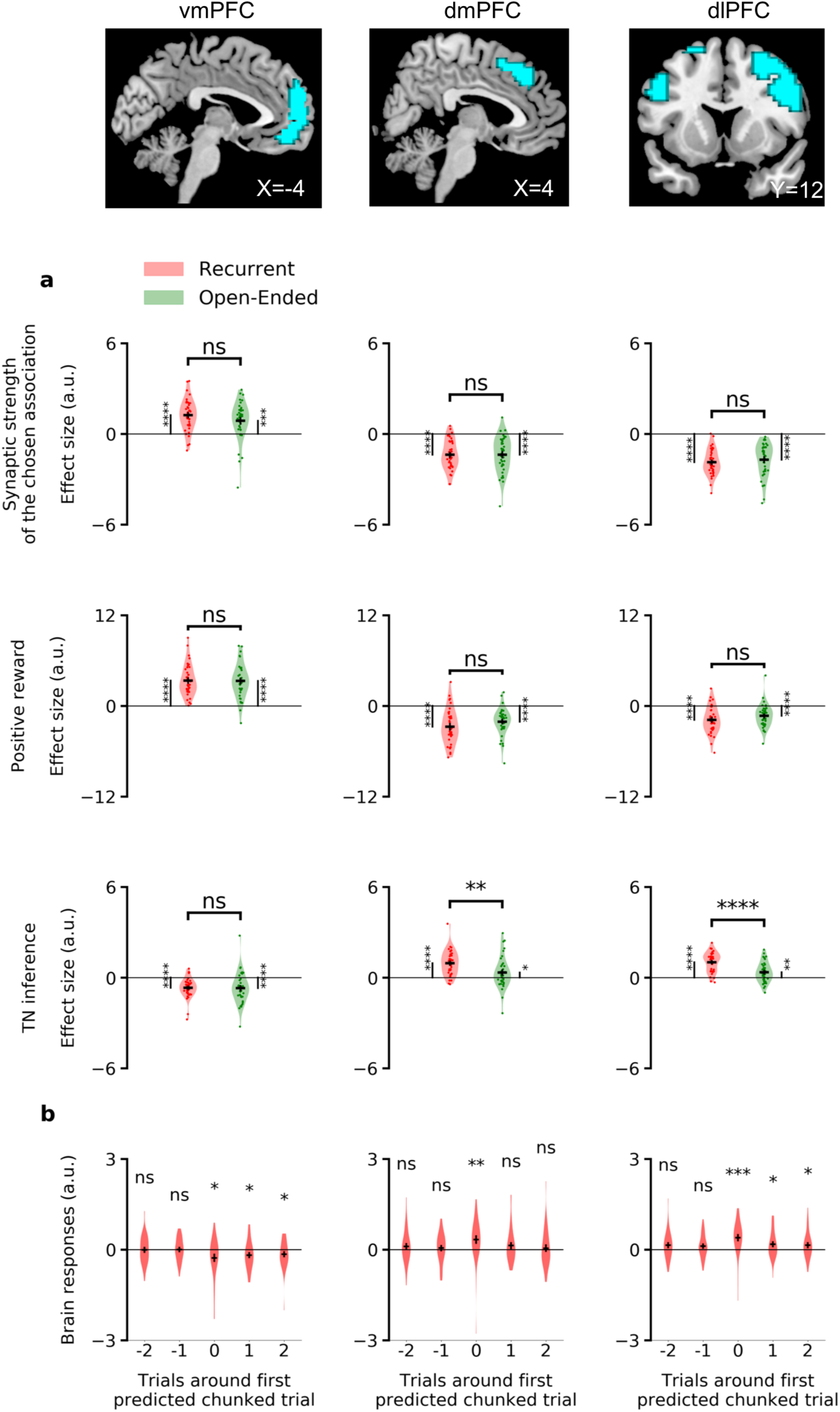
ROI analysis with the functional network defined from the ANOVA. The areas represented in blue are representing the regions identified in the previous analysis (Fig. 7) for the TN inference signal, using a significant threshold of FWE *p* = 0.05. **a,b,c**, They are tested at the onset feedback, thus independently, for the parametric modulators of the time series of *W*_*chosen*_, positive rewards, and the inference signal. **b**, Same GLM, in the recurrent session, but we replaced the parametric regressor of the task-set network inference signal by five sparser time-series constituted of only one trial per episode and the corresponding events shifted one or two trials preceding and following it. This trial was chosen as the first chunked trial of the model behavioral predictions (per episode, if it existed from sufficient learning, Fig. 6a,b,c). Effect sizes in arbitrary units for the recurrent and the open-ended session. Error bars correspond to the standard error of the mean over the 40 subjects. dlPFC: dorsolateral prefrontal cortex; dmPFC: dorsomedial prefrontal cortex; vmPFC: ventromedial prefrontal cortex

Because of the specific timing predicted by the inference signal, we pushed further this analysis by looking at the magnetic resonances responses at feedback onset, in these ROI, to a similar GLM where the inference signal had been replaced by the succession of 5 parametric regressors: an event-related regressor including only the first occurrence (per episode) of inference for a chunked trial (related to Fig. 6a,b,c), and the same event-related regressor shifted one or two trials preceding and following. This analysis is showed in Fig. 8b and confirmed that neural activity in these three ROI exhibits no response in the trials immediately preceding the first detected chunked trial. On the contrary, dorsomedial response is significant at the first detected chunked trial, and ventromedial and dorsolateral responses are significant from this trial onwards. The specific timing predicted by the model for this first detected chunked trial per episode is thus borne out both behaviorally (Fig. 6) and neurally (Fig. 8).

## Discussion

Task sets emerged in our model through unsupervised temporal chunking of stimulus-action associations encoded by mixed-selective neural populations. When repeated, a task-set could be retrieved from a single stimulus-action association by reactivation of the whole chunk. This retrieval then biased the downstream decision-making circuit through an inference signal. The model predicted finely timed, abrupt changes in behavioral responses in a task-set learning experiment [Collins and Koechlin, 2012; Donoso et al., 2014]. The retrieval of a task-set had both adaptive (reduction of exploration) and sometimes maladaptive effects (retrieval of an incorrect task-set) on the following trial performance. Our analysis of BOLD activity established a functional network engaging ventromedial, dorsomedial, and dorsolateral prefrontal cortex that correlated with the inference signal for task-set retrieval. The dorsal system was engaged preferentially in the situation where the retrieval of a task-set was used to improve performance.

### Biologically plausibility of the temporal chunking mechanism

The temporal chunking mechanism is implemented through simplified, but biologically plausible processes. The reward-dependent Hebbian learning and winner-take-all mechanisms of the associative network builds on previous studies in the field of conditional associative learning [Fusi et al., 2007; Soltani et al., 2017; Soltani and Wang, 2010; Wang, 2002; Wong and Wang, 2006]. Hebbian plasticity in the task-set network is slower, activity-mediated and unsupervised, creating task-sets as chunks of temporally contiguous events. The required mixed-selectivity can be obtained from randomly connected neurons receiving feed-forward inputs coming from sensory and motor areas and has been widely observed in the prefrontal cortex [Asaad et al., 1998; Genovesio et al., 2005; Rigotti et al., 2013; Wallis et al., 2001]. Moreover, learning at behavioral timescales can be generated through sustained neural activity and extended STDP [Brunel, 1996; Compte et al., 2000; Curtis and Lee, 2010; Drew and Abbott, 2006; Miller and Cohen, 2001; Murray et al., 2017; Rougier et al., 2005; Van Rossum et al., 2000]. Finally, the inference signal for task-set retrieval solves an exclusive-or problem, as different task-sets map from the same set of stimuli and actions. It can be implemented through random projections from an extra-pool of non-linear mixed selective cells both selective to the “internal” synaptic chunk (task-set) and to external events like the presentation of a stimulus [Fusi et al., 2016; Rigotti et al., 2013].

### Neural correlates of task-set retrieval

Consistent with the literature on the neural correlates of goal-directed behavior, we found that the inference signal predicting the retrieval of a task-set correlated specifically with BOLD activity in ventromedial, dorsomedial and dorsolateral prefrontal networks.

First, the associative network “prediction error” correlated with activity in striatum and ventromedial prefrontal cortex [Daw et al., 2006; Lebreton et al., 2009; Palminteri et al., 2015; Tanaka et al., 2004]. Ventromedial prefrontal cortex also correlated negatively with the inference signal, i.e. positively with the compatibility between encoding in the two subnetworks when a reward was received (as a prediction error-like signal). This is potentially in accordance with the role of ventromedial prefrontal cortex in monitoring the Bayesian actor reliability in this experiment [Donoso et al., 2014].

Second, dorsal prefrontal cortex was preferentially engaged when the model of the task (i.e., the task-sets) is useful and integrates into the behavioral policy (recurrent versus open-ended sessions, while controlling for trial perceived difficulty, as implemented by reaction times [Shenhav et al., 2013, 2014]). Dorsolateral prefrontal cortex is known to be specifically engaged for temporally integrating and organizing multimodal information [Duncan, 2001; Kim et al., 2008; Ma et al., 2014; Miller, 2000; O’Reilly, 2010; Sakai, 2008]. Previous work showed that neurons in the anterior cingulate cortex monitor the allocation and the intensity of control [Behrens et al., 2007; Dosenbach et al., 2006; Enel et al., 2016; Khamassi et al., 2013; Rushworth et al., 2007; Shenhav et al., 2013]. In this specific experiment, but through a Bayesian framework, the dorsal anterior cingulate cortex was shown to be specifically selective to switch-in events [Donoso et al., 2014].

### Stability/flexibility trade-off from the unsupervised temporal chunking mechanism

Our model builds on an attractor concretion mechanism [Rigotti et al., 2010b] while being simplified: we don’t make the hypothesis of the existence of fixed attractors. Instead, the synaptic weights are modified immediately from the start and continuously. Thus, the task-set network can learn from its own activity, combining prior statistical information to future learning, crucial in non-stationary problems. In order for this mechanism to be stable when learning concurrent task-sets, learning has to be slower as the representational complexity increases.

This mechanism also enables the encoding of a synaptic trace of any sequence of events, even incorrect, as a transition probability (weak but non-zero) between chunks or with an isolated neural population. The brain relies on estimates of uncertainty within and between task-sets [Collins and Koechlin, 2012; Courville et al., 2006; Donoso et al., 2014; Gershman and Niv, 2012; Kepecs and Mainen, 2012; Rushworth and Behrens, 2008; Soltani and Izquierdo, 2019; Yu and Dayan, 2005]. More specifically, Collins and Koechlin [Collins and Koechlin, 2012] have shown that the Bayesian likelihood (“reliability”) of each task-set in memory is evaluated. Inferences on the current and alternative task-sets have been found to occur in medial and lateral prefrontal cortices respectively [Donoso et al., 2014]. The coupling between these two tracks permits hypothesis testing for optimal behavior. Future work could investigate how an estimate of uncertainty (or reliability over task-sets) is retrieved from the synaptic weights of our model. This synaptic trace could then be used by the cholinergic and noradrenergic systems to modify the relative influence of top-down control (TN-like) versus bottom-up experience-dependent (AN-like), integration of information [Aston-Jones and Cohen, 2005; Sara, 2009], or to regulate the learning rates and the exploration parameters [Doya, 2002; Farashahi et al., 2017] responsible for the stability/flexibility trade-off of this temporal chunking mechanism.

### Temporal chunking in layers of mixed-selective cells is a plausible implementation of multi-step transition maps and generalization

Cognitive control and learning are linked and depend on the formation of hierarchical representations in the brain [Badre, 2008; Badre et al., 2009, 2010]. Plasticity and recurrence between mixed-selective cells in prefrontal cortex create a conjunctive code seeding a flexible “representational medium” [Duncan, 2001; Ma et al., 2014; Manohar et al., 2019; Miller, 2000]. Through simple Hebbian learning at decreasing timescales in decoupled layers of mixed-selective cells, momentarily stable chunks combining states, action, rewards, or more abstract “task-sets” can be encoded, as well as the transition statistics between them, to create hierarchically more complex chunks. A local change in the state transition function of the task, or during reward devaluation, will have the effect of flexibly depressing the newly unused synaptic link without relearning the whole transition rule. This “model-free” mechanism thus encodes the discounted occupancy of a state (or action, or any task-relevant feature), averaged over trajectories beginning from that state, leading to model-based behavior [Behrens et al., 2018; Russek et al., 2017] while also accounting for limited memory capacity [Blumenfeld et al., 2006; Nassar et al., 2018; Ostojic and Fusi, 2013].

This mechanism can also generalize. Augmenting our network with a generalization layer composed of neurons selective to the combination of three stimuli and three actions could produce faster learning of a new task-set by biasing lower cortical structures. In this simplistic scheme, generalization is a top-down, gating-like mechanism solving exclusive-or problems between layers of cells of decreasing complexity. Caching multi-steps transitions in a single value (model-free) or not (model-based) would then be equivalent to learning at slower timescales in an increasingly complex hierarchy of cortex layers [Murray et al., 2014].

## Acknowledgements

FB was funded by Ecole des Neurosciences de Paris Ile-de-France and Region Ile de France (DIM Cerveau & Pensee). SP is supported by an ATIP-Avenir grant (R16069JS) Collaborative Research in Computational Neuroscience ANR-NSF grant (ANR-16-NEUC-0004), the Programme Emergence(s) de la Ville de Paris, the Fondation Fyssen and the Fondation Schlumberger pour l’Education et la Recherche. SO is funded by the Ecole des Neurosciences de Paris Ile-de-France, the Programme Emergences of City of Paris, Agence Nationale de la Recherche grants ANR-16-CE37-0016-01 and ANR-17-ERC2-0005-01. This work was supported by the program “Investissements d’Avenir” launched by the French Government and implemented by the ANR, with the references ANR-10-LABX-0087 IEC and ANR11-IDEX-0001-02 PSL Research University.

## Competing Interests

The authors declare no competing interests.

## Materials and Methods

### Experimental procedures

#### The experimental task

We modeled a specific human experiment for concurrent task-set monitoring, previously reported in [Collins and Koechlin, 2012; Donoso et al., 2014]. The detail of the experimental procedures can be found in the original papers, here we provide only a summary. Data from [Collins and Koechlin, 2012] are called *Experiment 1*. Data from [Donoso et al., 2014] are called *Experiment 2*. The experimental designs are identical. *Experiment 1* is a behavioral experiment including 22 subjects. *Experiment 2* involves 40 subjects, with fMRI acquisition.

Subjects had to search for implicit associations between 3 digits and 4 letters by trial and error. In each trial, a visual stimulus (a digit on the screen in {1, 3, 5} or {2, 4, 6}) was presented to the subject (Fig. 1a). The subject had to take an action by pressing a letter on a keyboard in {*d, f, j, k*}. The outcome (reward or no reward) was announced with a visual and auditory feedback. A visual measure of the cumulative collected profit was displayed on the screen. For each trial of Experiment 1, the subject had 1.5 seconds to reply during the presentation of the stimulus. The average length of a trial was 2.9s. For Experiment 2, the mean of a trial was either 6s or 3.6s, whether BOLD activity is acquired or not. MRI trials were longer, introducing jitters at stimulus or reward onsets for signal decorrelation.

A *correct association* between the stimulus and the action led to positive reward with a probability 90%. An incorrect association between the stimulus and the action led to a null reward with a probability 90%. 10% of (pseudo-randomized) trials were misleading *noisy trials*, yielding to a positive reward for an incorrect association, and vice-versa. Thus a null feedback could be produced either by a behavioral error, by a change in correct associations, or by noise. The introduction of noisy trials prevented subjects from inferring a change in correct associations from a single unrewarded trial.

The correct set of responses to stimuli remained unchanged over a block of 36 to 54 trials. Such a block is called an *episode*. The transition from one episode to another is called an *episode switch* and was not explicitly cued. This transition imposes a change of correct responses from all stimuli, so a change of *task-set*. Task-sets were always non-overlapping from one episode to the other, i.e. each and every of the three stimulus-response associations differ after an episode switch. Within a given set, two stimuli were never associated with the same action.

An experimental *session* was a succession of 25 episodes for Experiment 1, and 24 episodes for Experiment 2. BOLD activity was acquired only during the 16 last episodes of Experiment 2.

In each experiment, subjects performed two distinct sessions: an open-ended and a recurrent session. In the *open-ended* session, task-sets were different in each episode, so there was no possibility for the subject to retrieve and reuse a formerly learned one. In the *recurrent* session, only 3 task-sets reoccurred across episodes (Fig. 1a), and subjects could reuse previously learned task-sets. Subjects were not informed about the distinction between the two sessions. The order of the sessions was changed between subjects to counteract for potential session-ordering effect. Different digits were used from one session to the other.

Having to manipulate at least 3 different task-sets was crucial: indeed when there are only 2 of them, the second one could be inferred from the first one by elimination. With 3 task-sets, and after an episode switch, some exploration is required to find the next mapping to use. More possible actions than the number of stimuli were used to avoid learning the third stimulus-response association by simple elimination when two associations were already known.

#### Debriefing

After each session, subjects performed a post-test debriefing. They were presented with 6 task-sets and rated them depending on their confidence in having seen them or not during the experiment. For the recurrent session, 3 out of the 6 task-sets were actually presented during the experiment. For the open-ended session, the 6 task-sets were all part of the experiment. From the debriefing of the recurrent session, subjects were classified in two different groups. *Exploiting subjects* ranked higher confidence for the 3 seen task-sets, compared to the 3 unseen task-sets. *Exploring* subjects, on the contrary, ranked at least 1 unseen task-set with more confidence than one of the 3 seen task-sets.

### Network model

The network model is based on [Rigotti et al., 2010b]. It is composed of two interacting subnetworks, the associative and task-set networks, illustrated in (Fig. 1d). In contrast to Rigotti et al. [2010b], we do not explicitly model temporal dynamics within a trial, but instead use simplified, instantaneous dynamics between populations replacing many mixed-selective neurons [Fusi et al., 2016]. Moreover the feedback from the task-set network to the associative network is implemented in a simplified manner. Full details of the model implementation are given below.

#### The associative network

The associative network (*AN*) is based on [Fusi et al., 2007]. This subnetwork implements in a simplified fashion the associations between input stimuli and output actions performed by a classical winner-take-all decision network [Brunel and Wang, 2001; Fusi et al., 2007; Wang, 2002].

The associative network is composed of neural populations selective to a single task-related aspect, either a stimulus or an action. Each population is either active or inactive in any trial, so that the activity is modeled as being binary. Any stimulus-selective neural population {*S*_*i*_}_*i*=1‥3_ projects excitatory synapses to all response-selective neural population {*A*_*j*_}_*i*=1‥4_. The corresponding synaptic strength is noted 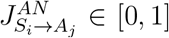. The behavioral output in response to a stimulus is determined based on these synaptic strengths, which moreover plastically change depending on the outcome of the trial (reward or no reward).

##### Action selection in the associative network

In any given trial, following the presentation of a stimulus *S*_*i*_, the associative network stochastically selects an action based on the strengths of the synapses from the population *S*_*i*_ = 1 to the populations that encode actions. Specifically, the action *A*_*j*_ is selected with the probability

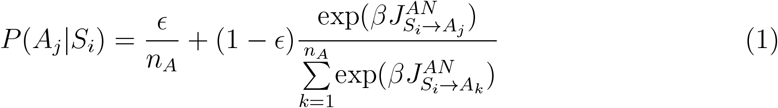

where *n*_*A*_ is the number of possible actions, 1/*β* is the strength of decision noise and *∊* accounts for the network’s internal estimate of *expected uncertainty* [Yu and Dayan, 2005]. The associative network therefore effectively implements a soft and noisy winner-take-all mechanism: all actions are equiprobable for high decision noise, whereas the probability of the action with the largest synaptic strength tends to 1 for low decision noise.

##### Synaptic plasticity in the associative network

Learning of the basic stimulus-action associations is implemented through plastic modifications of the synaptic strengths in the associative network. Following an action, the synaptic strengths are updated according to a reward-modulated, activity-dependent learning rule:

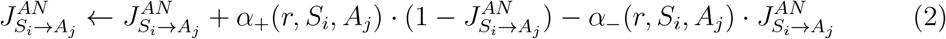

where *r* is the obtained reward (*r* = 0 or 1), and *α*_+_ and *α*_−_ are respectively the rates of potentiation and depression which depend on the reward as well as the activity of pre-and post-synaptic populations. Note that the update rule implements soft bounds on synaptic strengths, and ensures biological plausible saturation of neural activity, as well as forgetfulness [Amit and Fusi, 1994; Fusi, 2002; Fusi et al., 2007; Ostojic and Fusi, 2013].

The synaptic plasticity is local, so that only synapses corresponding to the active pre-synaptic population *S*_*i*_ are updated. Moreover, for simplicity, all non-zero potentiation and depression rates are equal and given by a parameter *α*. We therefore have

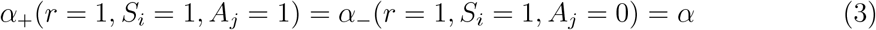

if the reward is positive, and

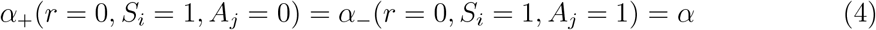

if the reward is null. All other potentiation and depression rates are zero.

The simplest implementation of the associative network therefore has three free parameters: the learning rate *α* and noise parameters *β* and *∊*. When fitting the model to human behavior, we have examined the possibility of adding complexity by distinguishing the learning rates corresponding to distinct reward and pre/post-synaptic events. The presented results on model fits, model dynamics and model predictions concerning the recurrent session in the present study are not modified by this extension.

### The task-set network

The task-set network (*TN*) is composed of mixed-selective neural populations, which are selective to conjunctions *S*_*i*_*A*_*j*_ of one stimulus and one action. As in the associative network, the activity of each population in the task-set network is represented as binary (either active or inactive). The task-set network is fully connected: any neural population *S*_*i*_*A*_*j*_ projects excitatory synapses to all other neural populations *S*_*k*_*A*_*l*_, with 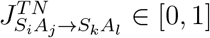. The strengths of these synapses are plastically updated, and determine the co-activation of populations in the task-set network. This co-activation effectively encodes task-sets. Full details of the model implementation are given below.

#### Activation of populations in the task-set network

At each trial, a stimulus *S*_*i*_ is presented and the associative network selects an action *A*_*j*_. In the task-set network, this leads to the activation of the population *S*_*i*_*A*_*j*_. Depending on the synaptic strengths, this may in turn lead to the co-activation of other populations in the task-set network. Specifically, if the synaptic strength 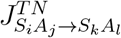 is greater than the parameter *g*_*I*_, the population *S*_*k*_*A*_*l*_ is activated. This step is iterated until no additional population gets activated. Here the parameter *g*_*I*_ represents an inhibitory threshold equivalent to a constant negative coupling between all populations in the task-set network, and implements in a simplified way a competition between excitatory neural populations through recurrent feedback inhibition.

These activation dynamics are assumed to be fast on the timescale of a trial, and therefore implemented as instantaneous.

#### Synaptic plasticity in the task-set network

The synapses in the task-set network are updated following an unsupervised, Hebbian plasticity rule. This update is driven by the sequential activation of neural populations in the task-set network, and thus by the associative network dynamics (Fig. 1d).

When two populations in the task-set network are activated on two consecutive trials, the synapses connecting them are potentiated. Noting *S*_*t*_*A*_*t*_ the task-set network neural population active at trial *t*, and *S*_*t*+1_*A*_*t*+1_ the neural population active at trial *t* + 1, this potentiation is given by

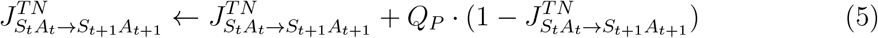

where the parameter *Q*_*P*_ represents the learning rate for potentiation.

At each trial, all the synapses from the active neural population *S*_*t*_*A*_*t*_ = 1 are depressed (*pre-activated depression* [Ostojic and Fusi, 2013]), implementing an effective homeostatic control. This depression is given by

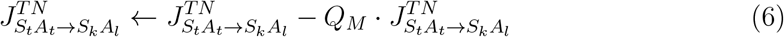

where *Q*_*M*_ is the rate of depression.

The ratio *Q*_*p*_/*Q*_*m*_ between the potentiation and the depression rates determines the asymptotic values of the synaptic strengths [Ostojic and Fusi, 2013]. To produce co-activation of populations in the task-set network and therefore the learning of task-sets, this asymptotic value needs to be higher than the inhibition threshold *g*_*I*_. To avoid any redundancy between the parameters we fixed *g*_*I*_ to 0.5 and 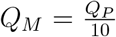, so that *Q*_*p*_ is the only free parameter.

#### Interaction between associative network and task-set network

To implement the effect of the task-set network dynamics and learning on the output of the network, the pattern of activity in the task-set network needs to influence the activation in the associative network. In our model this interaction is implemented in a simplified fashion.

If the previous trial was rewarded, the activation in the associative network on the next trial is biased towards the stimulus-action combinations that correspond to activated populations in the task-set network. This effective *inference signal* is implemented by modulating the strength of synapses in the associative network. Specifically, If *r* = 1 and *S*_*k*_*A*_*l*_ = 1 in the task-set network on the previous trial, the synaptic strength from *S*_*k*_ to *A*_*l*_ in the associative network is increased according to

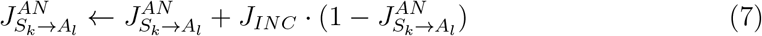

where the parameter *J*_*INC*_ specifies the strength of the inference signal.

The strength of the inference signal thus corresponds to the discrepancy between the task-set network prediction and the associative network encoding when a reward is received (or its negative counterpart, the compatibility).

### Model fitting to behavioral data and simulation

The model without task-set inference has 3 free parameters, and the model with task-set inference has 5 free parameters. The parameter set is composed of the associative network learning rate *α*, the task-set network learning rate *Q*_*P*_, the parameters of the soft and noisy winner-take-all mechanism (decision noise 1/*β* and uncertainty ∊), and the inference strength *J*_*INC*_ from task-set network to associative network connectivity. Both models were fitted to behavioral data using the standard maximum likelihood estimation (MLE). We provide the model with the subject’s set of actions and we define the model *performance on a trial* as the model’s likelihood of observing the subject’ response at the next trial. The Bayesian information criterion (Fig. 5a) uses the likelihood computed by MLE but introduces a penalty term for model complexity depending on the size of the observed sample. We also compared the AIC (Akaike information criterion) for both models and reached identical conclusions. A larger log-likelihood and lower BIC and AIC correspond to best model fits. We combined a grid search on initial parameters values with a gradient descent algorithm from the SciPy optimization toolbox. Parameters were estimated subject by subject.

On average over subjects (see Fig. S1), the learning rate in the associative network (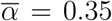, *σ* = 0.0073) is twice the learning rate in the task-set network (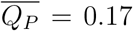, *σ* = 0.0070). The inference strength from the task-set network to the associative network is high in the recurrent session (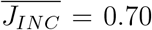, *σ* = 0.037), and its value is significantly lower in the open-ended session (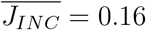, *σ* = 0.0062, see Fig. 5b).

We also compared model simulations ex post (model recovery [Palminteri et al., 2017]), with and without task-set inference. In a simulation, the model’s actions are random depending on the trial by trial probability set computed from the winner-take-all mechanism. The *model performance* is now the probability predicted by the model for the correct action, at each trial. Model simulation reproduce model fits and data, which ensures that we are not overfitting subjects’ data.

### fMRI whole brain analysis

The model-based fMRI analysis was performed with SPM 12. The detail of the data acquisition can be found in [Donoso et al., 2014].

All parametric modulators are z-scored to ensure between regressor and between subjects commensurability [Lebreton and Palminteri, 2016]. For each onset, they are orthogonalized to avoid taking into account their shared variance. fMRI data were analyzed using an event-related General Linear Model. Regressors of non-interest include subjects’ lapses, post-pause trials at the beginning of each scanning run, and movement parameters. Event-related regressors of interest modeled separately the onset decision (stimulus presentation timing, covering the decision time window) and the onset feedback (outcome presentation timing). The regressors are based on the best fitting parameters at the subject level.

At the onset decision, the regressor includes orthogonalized parametric modulations following this order:

- The first modulator is the time-series of reaction times, an index of trial difficulty and a specific motor preparation-related activity.
- The second modulator is the associative network synaptic strength from the presented stimulus selective neural population to the chosen action selective neural population. We call this parameter *W_chosen_* and it is also an index of trial difficulty.

At the onset feedback, the regressor includes orthogonalized parametric modulations following this order:

- The first parametric modulator is *W*_*chosen*_. This control ensures that the correlations observed are not simply caused by the monitoring of the certainty on the chosen association (prediction error) or else the trial difficulty.
- The second parametric modulator is the time series of positive rewards.
- The third parametric modulator is the trial-by-trial average value of the inference signal from the task-set network to the associative network. It is thus the average, over the number of connections implicated, of the task-set network inference on the update of associative network synaptic weights. We call it *TN inference*.

All the mentioned time series are convolved with the hemodynamic response function to account for the hemodynamic lag effect.

The subject by subject statistical maps are combined to make generalizable inferences about the population. We use a *random effect analysis* approach [Holmes and Friston, 1998]. We identify activations using a significance threshold set to *p* = 0.05 (familywise error FWE corrected for multiple comparison over the whole brain).

For conciseness, and because and mixed-selectivity has been found in prefrontal cortex [Fusi et al., 2016], we do not report posterior activations (parietal, temporal and occipital lobes).

Note that we did a preliminary control analysis using the link between the associative network and Q-learning [Watkins and Dayan, 1992] by searching for any correlation between BOLD activity and the prediction error, i.e. the difference between the perceived outcome and the associative network synaptic strength of the trial-by-trial chosen association. As expected from previous studies [Daw et al., 2006; Kim et al., 2006; Lebreton et al., 2009; O’Doherty et al., 2004; Tanaka et al., 2004], we found ventromedial prefrontal cortex and striatal activity to correlate positively in the recurrent and in the open-ended session. The MNI peak coordinates and number of voxels in the cluster were respectively [−12, 56, 20], *T* = 11.2, 901 voxels in the recurrent session, and [−12, 8, −12], *T* = 14.3 and 1419 voxels in the open-ended session.

We first investigated neural correlates of the trial-by-trial synaptic strength of chosen association in the associative network, at the onset decision (*W*_*chosen*_), and the neural correlates of the trial-by-trial inference signal strength from the task-set network to the associative network, at the onset feedback. Results are shown in Fig. S5.

### One-way ANOVA and second control ROI analysis

In order to test the hypothesis of a specific effect of task-set retrieval, we extract the betas from medial and lateral prefrontal nodes, and compare them from the two conditions: the recurrent and the open-ended session. This comparison is valid as soon as the region of interest is selected independently from the statistical maps of betas [Poldrack, 2007], i.e. the selected ROI need to be based on a different contrast that the one currently studied. We defined a functional network by the co-activations in both sessions, for the trial-by-trial task-set network inference signal to the associative network (ANOVA REC+OE for dorsomedial and dorsolateral prefrontal cortex, ANOVA -REC-OE for ventromedial prefrontal cortex, FWE 0.05, Table 7), which thus did not promote differences. Our ROI of ventromedial, dorsomedial and dorsolateral prefrontal cortex were selected from the obtained thresholded maps (FWE *p* = 0.05) from this ANOVA analysis, and were used to test differences between REC and OE.

We further controlled our results by running other independent ROI analysis using:

- the Stanford Functional Imaging in Neuropsychiatric Disorders Lab [Shirer et al., 2012]
- the Glasser parcellation [Glasser et al., 2016]
- the Neurosynth meta-analysis [Yarkoni et al., 2011]
- the WFU PickAtlas [Lancaster et al., 2000; Lancaster, 1997; Maldjian et al., 2003] Results are shown in Fig. S6.

### Software

All simulations were done with Python 2.7 (using numpy and scipy, and the scikit-learn package [Pedregosa et al., 2011]. The fMRI analysis was done with Matlab and SPM12 [Ashburner et al., 2014].

## Supplemental Figures

**Figure S1:**
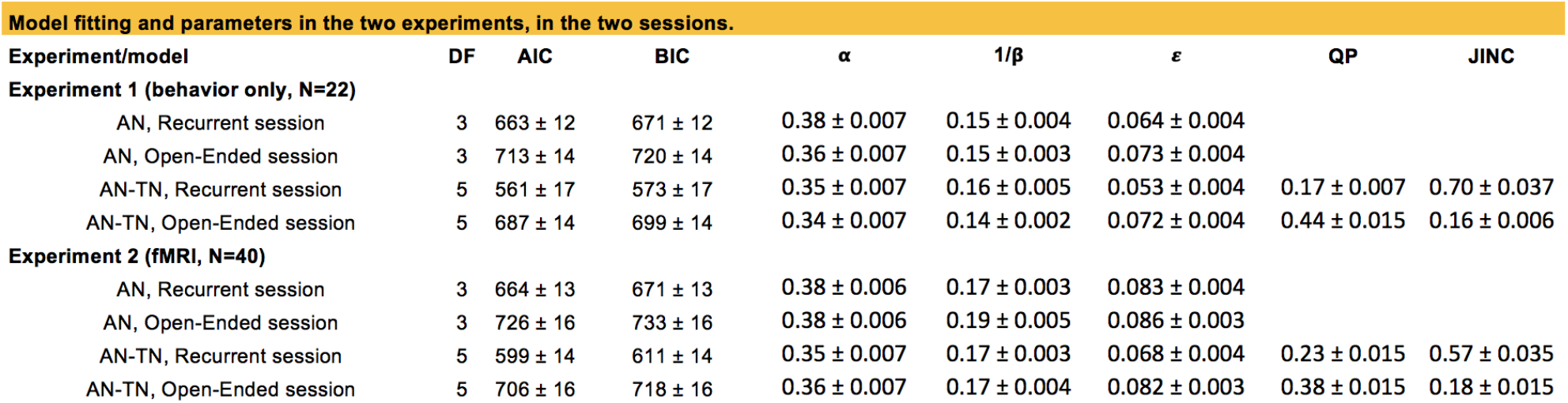
The table summarizes the full network (AN-TN, with inference) and the associative network alone (AN, without inference) models fitting performances and average parameters. Related to Fig. 5. DF, degrees of freedom; AIC, Akaike information criterion; BIC, Bayesian information criterion; *α*, learning rate in the AN; 1/*β*, decision noise; ∊, uncertainty; *Q*_*P*_, learning rate in the TN; *J*_*INC*_, inference strength. All are expressed as mean ± s.e.m.

**Figure S2:**
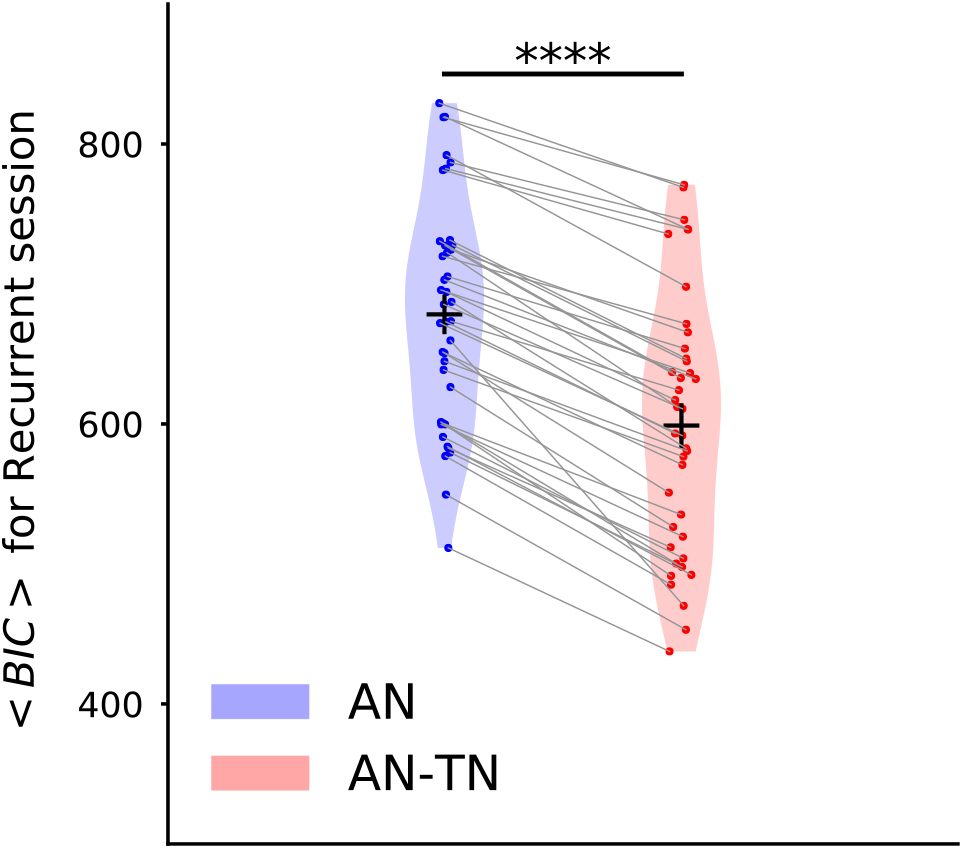
Model comparison for the recurrent session, for Experiment 2. Related to Fig. 5. Bayesian Information Criterion, computed as in [Bishop, 2007], for the models with and without task-set inference. The model provides a significantly better fit with inference than without (*p* = 9.1 · 10^−5^, *t* = 4.1).

**Figure S3:**
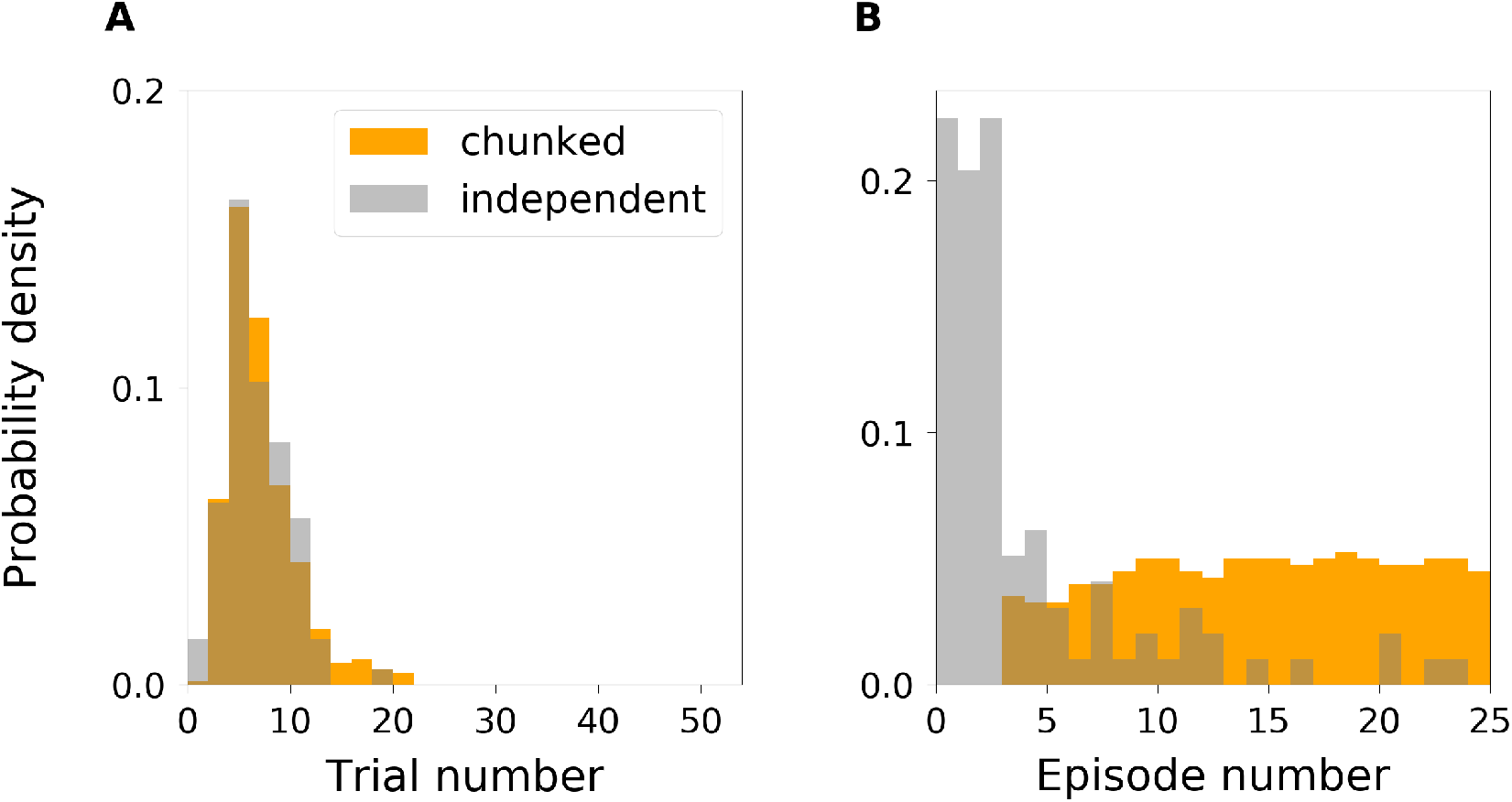
Task-set retrieval prediction. Experiment 1. Related to Fig. 6a,b,c. **a**, Distributions of trial numbering for the two categories of trials, chunked and independent. The distributions are not significantly different (a Kolmogorov-Smirnov test gives *ks* = 0.085, *p* = 0.62). **b**, Distributions of episode numbering for the two categories of trials, chunked and independent. We consider only one trial per episode. Generally, *independent* trials are from early episodes, and *chunked* trials are from late episodes, consistently with the expected learning progress.

**Figure S4:**
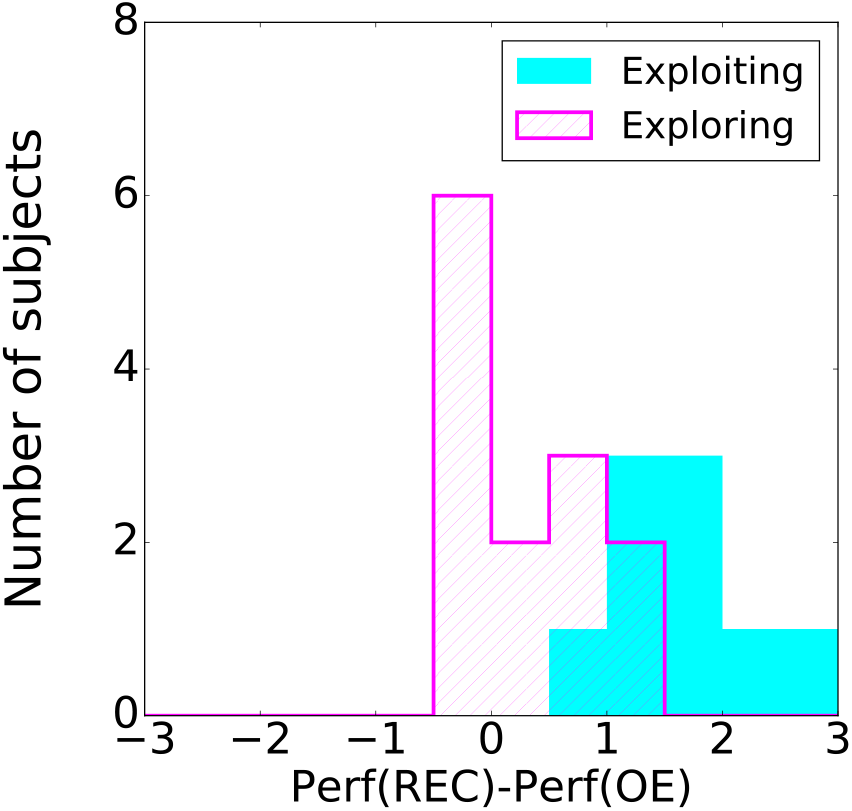
Experiment 1 - Histograms over subjects of the difference of performance after 5 first consecutive correct trials, between the recurrent session and the open-ended session. Related to Fig. 6. The classification of subjects is based on the model prediction. The difference between the two distributions is statistically significant (a Kolmogorov-Smirnov test gives *p* = 3 *·* 10^−4^)

**Figure S5:**
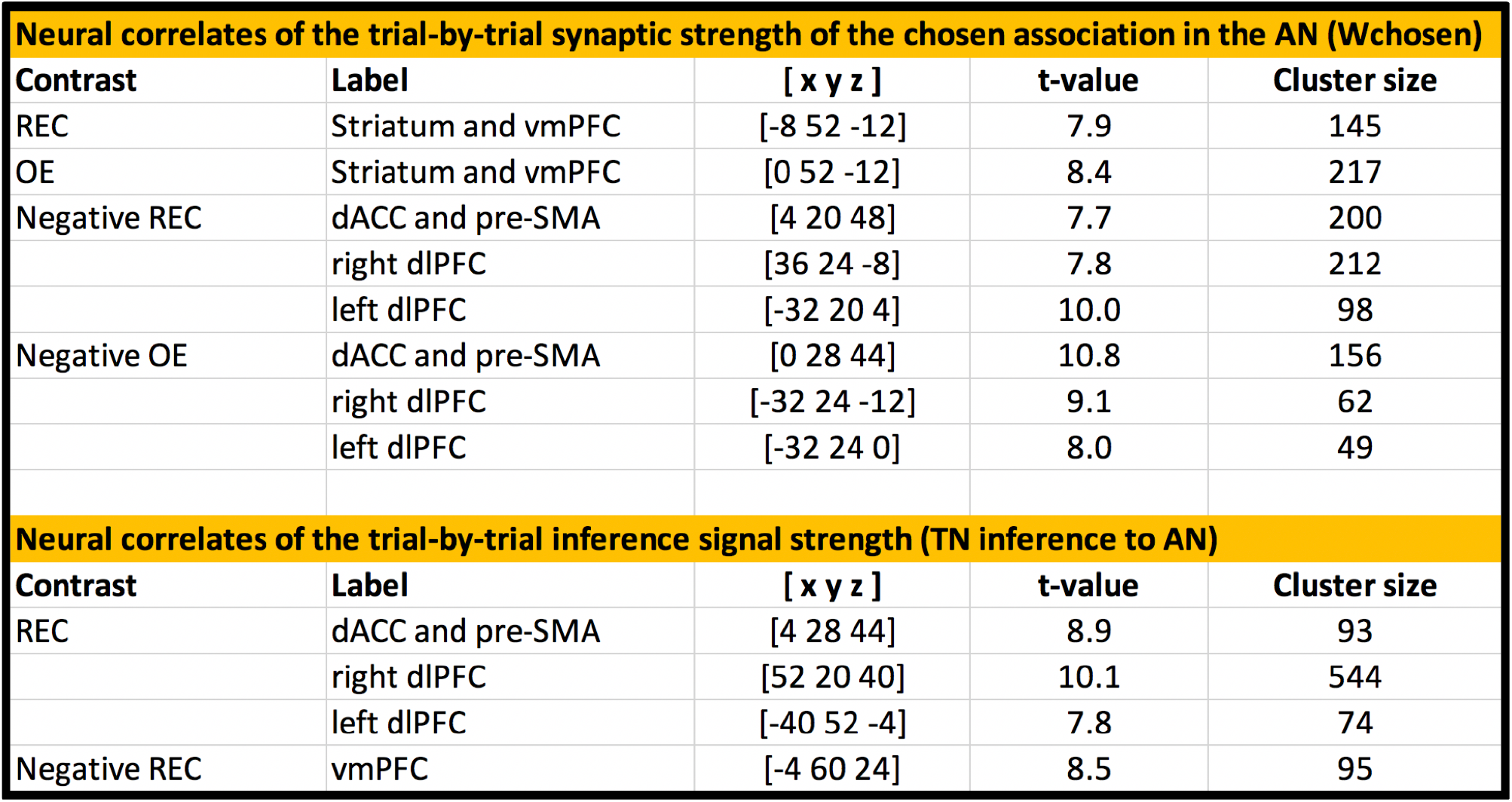
Neural correlates of the synaptic strength in the associative network, and of the inference from the task-set network to the associative network. top, Activations (FWE *p* = 0.05) from the parametric modulator corresponding to the synaptic strength of the chosen association in the associative network, *W*_*chosen*_, at the onset decision. **bottom**, Activations (FWE *p* = 0.05) from the parametric modulator corresponding to the inference from the task-set network to the associative network, at the onset feedback. No activation (FWE *p* = 0.05) was found in the open-ended session. dlPFC: dorsolateral prefrontal cortex; dmPFC: dorsomedial prefrontal cortex; vmPFC: ventromedial prefrontal cortex; [x y z] are MNI coordinates; REC: Recurrent session; OE: Open-Ended session.

**Figure S6:**
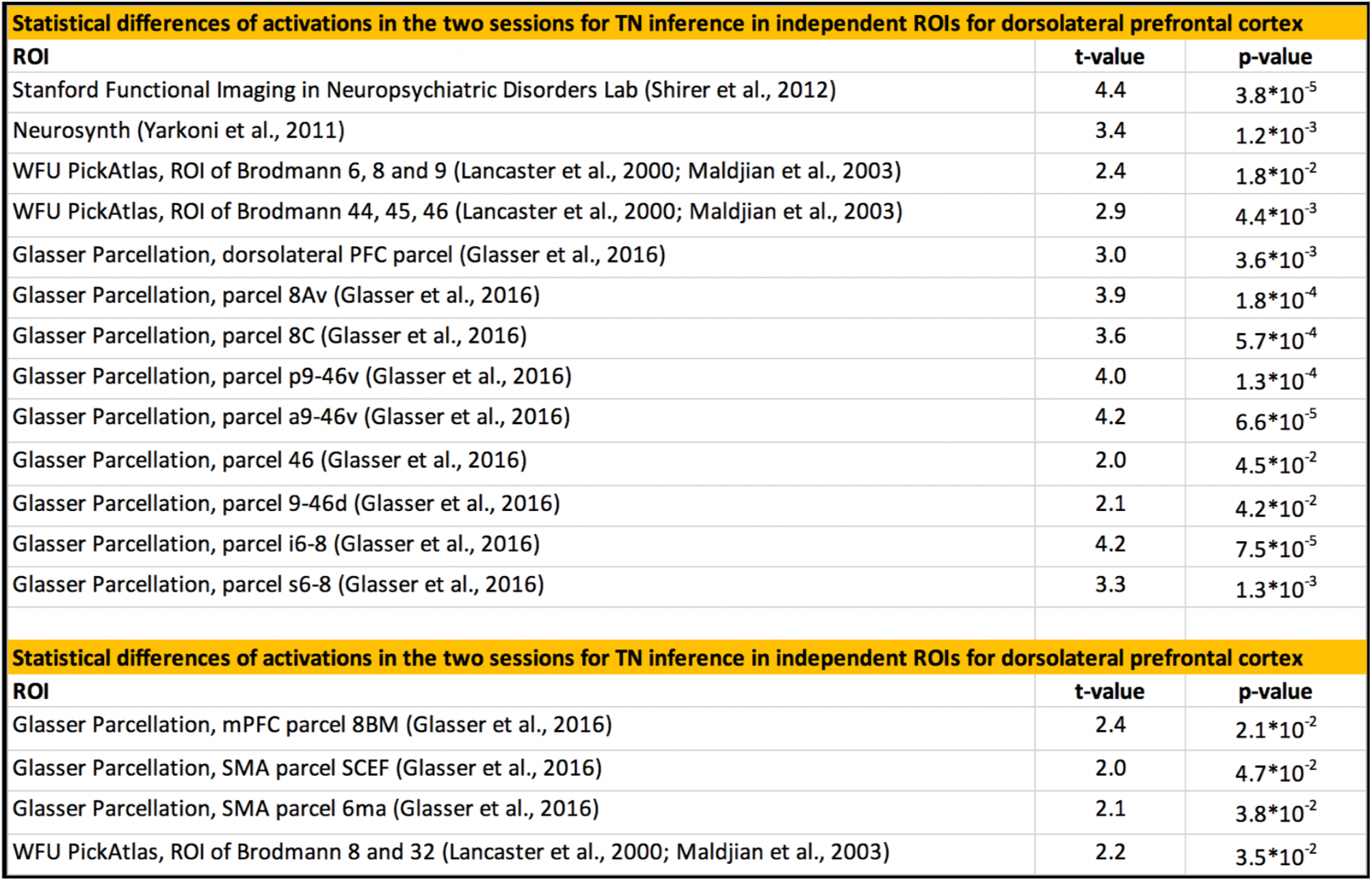
Control independent ROI analysis: neural correlates of the inference signal from the task-set network to the associative network, at the onset feedback. top, Statistical difference between activations in the recurrent session and in the open-ended session, in independent ROIs for dorsolateral prefrontal cortex. **bottom**, Statistical difference between activations in the recurrent session and in the open-ended session, in independent ROIs for dorsomedial prefrontal cortex. dlPFC: dorsolateral prefrontal cortex; dmPFC: dorsomedial prefrontal cortex; vmPFC: ventromedial prefrontal cortex; [x y z] are MNI coordinates.

